# Model-directed generation of CRISPR-Cas13a guide RNAs designs artificial sequences that improve nucleic acid detection

**DOI:** 10.1101/2023.09.20.557569

**Authors:** Sreekar Mantena, Priya P. Pillai, Brittany A. Petros, Nicole L. Welch, Cameron Myhrvold, Pardis C. Sabeti, Hayden C. Metsky

**Affiliations:** Broad Institute of MIT and Harvard, Cambridge, MA, USA; Department of Statistics, Harvard University, Cambridge, MA, USA; Department of Molecular and Cellular Biology, Harvard University, Cambridge, MA, USA; Division of Health Sciences and Technology, Harvard Medical School and Massachusetts Institute of Technology, Cambridge, MA, USA; Harvard/Massachusetts Institute of Technology, MD-PhD Program, Boston, MA, USA; Department of Systems Biology, Harvard Medical School, Boston, MA, USA; Department of Molecular Biology, Princeton University, Princeton, NJ, USA; Howard Hughes Medical Institute, Chevy Chase, MD, USA; Department of Organismic and Evolutionary Biology, Harvard University, Cambridge, MA, USA; Department of Immunology and Infectious Diseases, Harvard T.H. Chan School of Public Health, Boston, MA, USA

## Abstract

Generating maximally-fit biological sequences has the potential to transform CRISPR guide RNA design as it has other areas of biomedicine. Here, we introduce model-directed exploration algorithms (MEAs) for designing maximally-fit, artificial CRISPR-Cas13a guides—with multiple mismatches to any natural sequence—that are tailored for desired properties around nucleic acid diagnostics. We find that MEA-designed guides offer more sensitive detection of diverse pathogens and discrimination of pathogen variants compared to guides derived directly from natural sequences, and illuminate interpretable design principles that broaden Cas13a targeting.

## Main text

Machine learning methods are transforming the design of biological sequences^1–5^. Generative methods, in particular, have been extensively applied to designing proteins^6–12^, yet other uses of generative methods, such as designing CRISPR guide RNA sequences, remain relatively underexplored. Current techniques^13–22^ for guide RNA design select sequences directly from those in nature or from a simple function of natural sequences, e.g., a consensus, according to machine-learned discriminative models or heuristic rules. However, natural sequences may not perform the best in every application. Generative design methods could yield artificial guide RNA sequences—that is, guides that differ from any natural sequence, sometimes by 3 or more mismatches—and achieve superior performance on particular tasks. To our knowledge, such techniques have not been applied to design CRISPR guides.

Pathogen detection has been one promising use of CRISPR-based technologies. The extensive genomic diversity of pathogens motivates using generative methods to design guides that can optimally detect them. For instance, if our goal were to detect all forms of a genetically diverse pathogen (“multi-target detection”), and there are several polymorphisms in an otherwise efficacious target region, an artificial guide could more sensitively detect all combinations of polymorphisms than any guide that fully matches one particular combination (Fig. 1a). This would enable us to achieve improved diagnostic sensitivity. Alternatively, if our goal were to distinguish between two highly similar targets (“variant identification”), even those that differ by a single nucleotide polymorphism (SNP), an artificial guide with optimally-positioned mismatches could achieve greater specificity than a guide identical to one target (Fig. 1b). This would enable us to more accurately identify mutation(s) in a pathogen or discriminate between pathogen lineages. Previous work^23, 24^ has described heuristic rules to introduce handcrafted specificity-enhancing mismatches into CRISPR guides, but these strategies are bespoke and are limited by the degree of variation they can distinguish.

**Figure 1.**
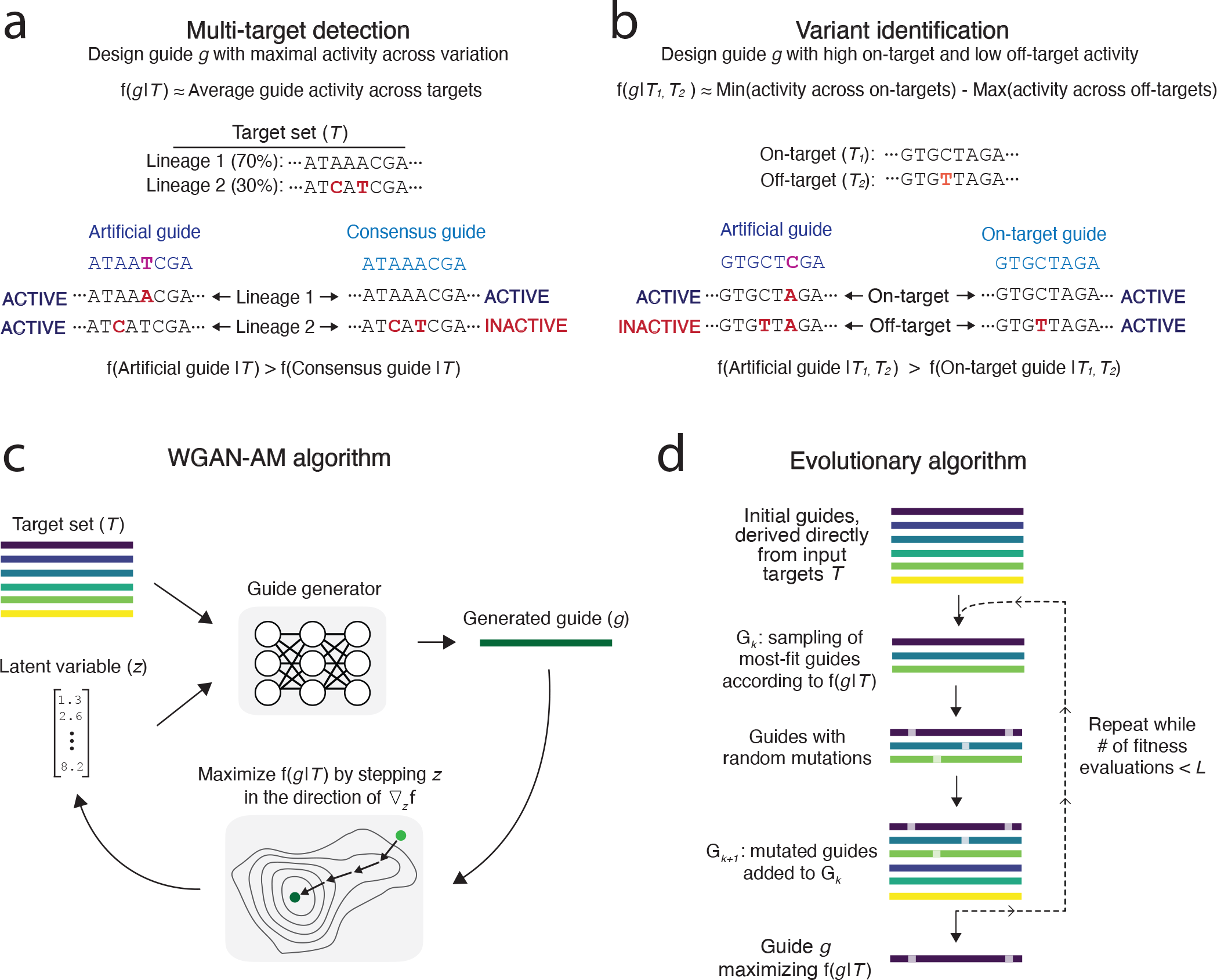
Designing optimal guides for two diagnostic objectives using model-directed exploration algorithms. **(a)** Artificial guide sequences — i.e., those that differ from any naturally observed sequence — can increase detection across sequence variation. In the example target set shown, there are two lineages. The consensus of the targets has two mismatches with lineage 2 (in red), which could reduce enzymatic activity and thus sensitivity for that lineage. The artificial guide sequence, which has only one mismatch with lineage 1 and one with lineage 2, could yield superior performance, since Cas13a better tolerates a single mismatch than two mismatches in close proximity. **(b)** Artificial guide sequences can increase the specificity of a guide designed to discriminate one target from another. A baseline approach is to design a guide whose sequence is directly derived from the on-target sequence. Here, the baseline guide has only one mismatch against the off-target sequence, and thus has substantial off-target activity and achieves poor specificity. An artificial guide sequence that has an additional mismatch introduced could have a double mismatch to the off-target sequence, enabling lower off-target activity and more robust specificity. **(c)** WGAN-AM algorithm for guide design. The generative model, G(*z* | *T*), generates active guide sequences conditional on a given target set *T* . A latent variable *z* modulates the generator’s output, allowing for different guide sequences. To generate optimal guides, the algorithm starts at a random value of *z*, computes the fitness of the generated guide f(*g* | *T*), and adjusts *z* in the direction of ∇_*z*_ f(*g* | *T*). **(d)** Evolutionary algorithm for guide design. The algorithm initializes a population of candidate guides, G_0_, by extracting the guide-length subsequences at the targeted genomic site. To form the population G_*k*_ in each generation *k, n* parent guides are sampled from the population G_*k*−1_ with probability proportional to their fitness f(*g* | *T*). The *n* guides are randomly mutated, and the resulting mutated child guides are added to the population. This process is repeated until *L*, the limit on the number of guides whose fitness has been evaluated, is reached.

Designing artificial guides—here, 28-nt Cas13a guide RNA spacer sequences—necessitates exploring a vast high-dimensional space of RNA sequences. Each guide has a fitness according to the task at hand, such as multi-target detection (Fig. 1a) or variant identification (Fig. 1b). Selecting a natural genomic sequence, as is typically done, amounts to exploring the very limited set of observed sequences at a site; more thoroughly exploring the sequence space could yield artificial guides with superior performance. Highly fit artificial guides should be nearby in sequence space to the complements of their targets, but this region, with ∼10^7^–10^9^ sequences (Supplementary Note 1), is sufficiently large to motivate the development of algorithms that efficiently search the sequence space.

Here, we develop model-directed exploration algorithms (MEAs) for CRISPR-Cas13a guide RNA design. MEAs combine a machine-learned model—trained to predict the enzymatic activity that results from a guide-target interaction—with search algorithms to explore a fitness landscape of guides. Our model is a convolutional neural network (CNN) that we previously published^22^. It predicts Cas13a’s enzymatic activity, a proxy of a given guide’s sensitivity for detecting a given target. We evaluate two search algorithms: (i) a generative model that employs a conditional Wasserstein Generative Adversarial Network (WGAN) and activation maximization (AM) to explore a continuous latent space of sequences^25^ (Fig. 1c and Supplementary Fig. 1); (ii) an evolutionary algorithm that performs iterative rounds of random mutation, fitness evaluation, and selection to explore the fitness landscape in discrete steps (Fig. 1d and Supplementary Fig. 2). Importantly, our model-directed exploration process—including the predictive model, search algorithms, and fitness functions—is conditioned on user-provided target sequences, making it broadly applicable to any input targets rather than being restricted to specific genomic sites.

First, we applied our methods to design guides, for use with Cas13a-based detection, that are sensitive across a pathogen’s genomic variation, i.e., for multi-target detection. The fitness function that we maximize is the average predicted activity across sequenced genomes (Fig. 1a, Methods). We focus on five RNA viruses, chosen for their extensive diversity (enterovirus B, Lassa, dengue) or public health relevance (influenza A, SARS-CoV-2). We employed the MEAs to design guides and benchmarked them against algorithms representing current, state-of-the-art design approaches (Methods). These include a simple, commonly-used algorithm that complements the consensus sequence at the target site (“consensus”) and a more advanced algorithm that we term “model-based choice” (MBC), which reflects a method we previously developed^22^. MBC employs clustering to compute a candidate set of guides and selects the guide with maximal fitness based on our model’s predictions (Methods).

We computationally evaluated the multi-target detection performance of the different design methods across these five pathogens using the same predictive model of guide activity (Methods). MEA-designed guides were predicted to detect more diversity than guides designed by baseline methods across nearly all genomic regions of all five pathogens. In several regions within dengue, Lassa, and influenza A viruses, the MEA-designed guides were predicted to detect 10–20% more sequence variation (Fig. 2a and Supplementary Fig. 3). The MEA-designed guides also achieved a higher mean fitness genome-wide (Fig. 2b and Supplementary Fig. 4). At most sites, the guides generated by the MEAs are several mismatches away from any sequenced genomes, suggesting that the MEAs’ ability to search fitness landscapes and generate artificial sequences contributes to their superior performance (Fig. 2c and Supplementary Fig. 5).

**Figure 2.**
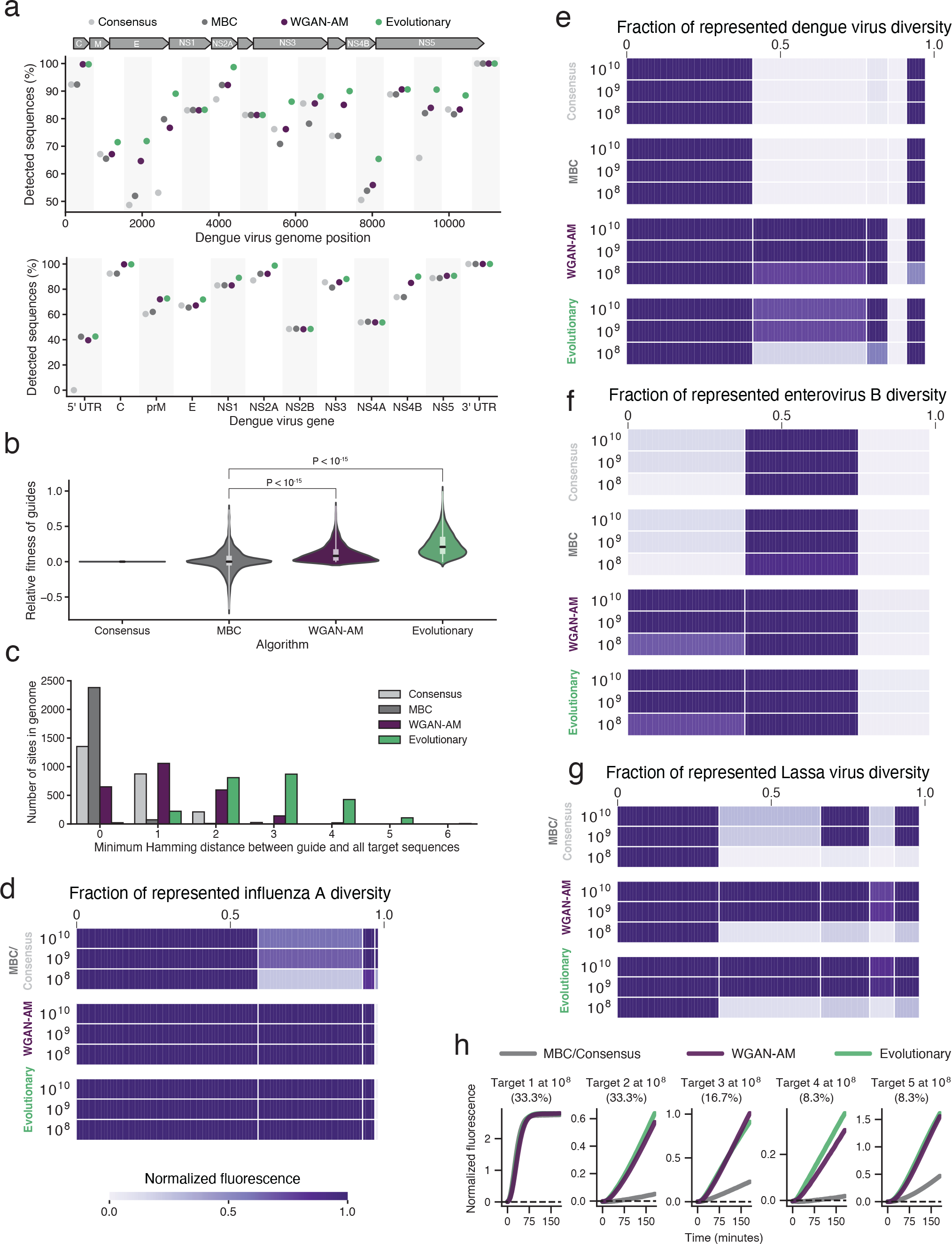
MEAs design guides that are maximally sensitive across genomic variation. **(a)** Proportion of dengue virus genomes detected by the diagnostic guides designed with model-based exploration algorithms (WGAN-AM and evolutionary) and baseline methods (model-based choice (MBC) and consensus). Top, within fixed-size windows along the genome; bottom, within genes and untranslated regions (UTRs). The MBC guides were designed by computing a ground set of sequences and employing the predictive model to select the sequence with the highest fitness. The consensus guides were designed by computing the consensus of the multiple sequence alignment at the targeted genomic site. A guide is considered to detect a target if it meets the criteria described in the Methods. **(b)** Relative fitness of the guides designed by MBC, WGAN-AM, and evolutionary algorithms at sites in the dengue virus genome. The relative fitness is the difference between the fitness of the labeled algorithm’s guide and the fitness of the consensus guide. The distribution is across targetable genomic sites (see Methods). *P* values were computed using one-sided Wilcoxon signed-rank tests. **(c)** Minimum Hamming distance between the guides designed by each algorithm and all target sequences at a given genomic site. Distribution is shown across targetable genomic sites. **(d–g)** Normalized fluorescence for guides detecting representative targets of genomic sites in influenza A (d), dengue virus (e), enterovirus B (f), and Lassa virus (g) at one hour. Each column represents a target and has width proportional to the percentage of sequence diversity it represents. Each row is a concentration of the target sequence in copies/μL **(h)** Normalized fluorescence, over the course of the reaction, when detecting the five representative targets of Lassa virus at 10^8^ copies/μL. Parentheticals indicate the percentage of all genomes represented by the target. The MBC-designed guide and consensus guide were identical at this site, and are represented by ‘MBC/consensus’.

Computational predictions are inherently limited, so we set out to experimentally benchmark the multi-target detection performance of the MEA-designed guides. We used a random sampling strategy (Methods) to select three representative genomic sites from each of the five viruses we considered (15 sites in total) and employed both the MEAs and baseline methods to design guides for these sites. We tested these guides against multiple synthetic targets that represent the genomic diversity of each virus (Methods), measuring the fluorescence readout over time. Cas13a is activated through guide-mediated recognition of a target; once active, it engages in collateral cleavage of a fluorescent reporter to generate the fluorescent readout, with greater fluorescent signal indicating stronger guide-target affinity at a fixed target concentration. For all but one of the 15 tested sites, the MEA-designed guides achieved higher activity across sequence diversity than the baseline guides (Fig. 2d–g, Supplementary Fig. 6, and Supplementary Fig. 7) and often enabled a lower limit of detection (LoD) as well (Fig. 2h and Supplementary Fig. 6). The MEAs provide the greatest advantage in highly diverse viruses, e.g., Lassa virus; this advantage is less pronounced at more conserved sites from less diverse viruses, e.g., SARS-CoV-2 (Supplementary Fig. 3 and Supplementary Fig. 8).

We next applied the MEAs to our second diagnostic objective: to design guides that optimally identify specific SNPs or distinguish between pathogen lineages, i.e., variant identification (Fig. 1b). This objective can be conceptualized as designing a guide that achieves high activity on an on-target sequence and minimal activity on an off-target sequence; the fitness function is approximately the difference between these activities (Fig. 3a, Methods).

**Figure 3.**
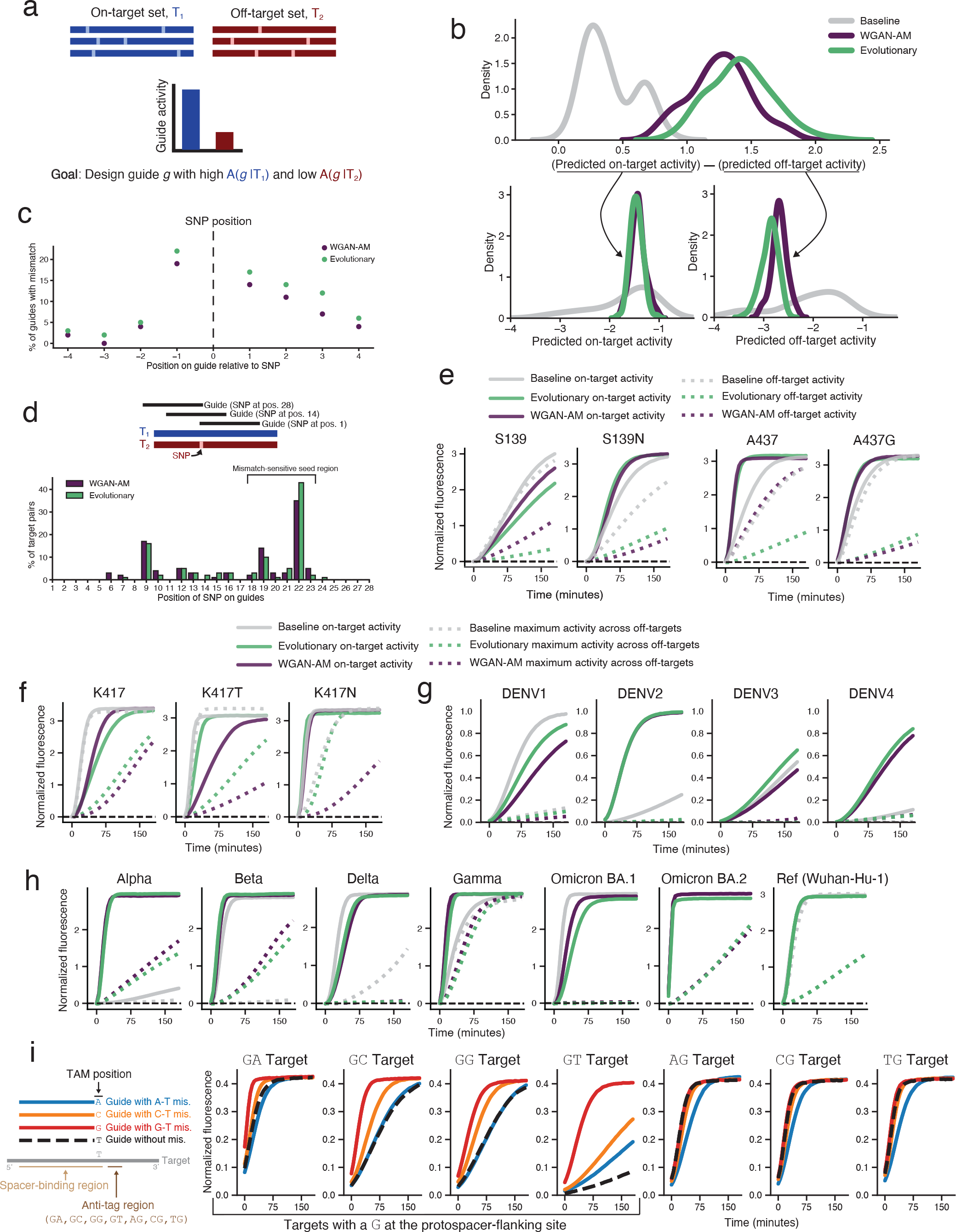
MEAs design optimal guides for variant identification and propose a new mechanistic guide design principle. **(a)** The objective of the variant identification task is to design a guide sequence that has maximal activity across a set of on-target sequences *T*_1_ and minimal activity across a set of off-target sequences *T*_2_. **(b–d)** Design features of 28-nt guides that differentiate between 100 pairs of synthetic targets such that, in each pair, *T*_2_ has a single nucleotide difference (1 SNP) with *T*_1_. For each pair of targets, we applied the MEAs to design guides for each of the 28 possible SNP placements along the guide, and chose the guide with the highest fitness among the 28 as best able to differentiate the two targets. **(b)** Benchmarking predicted activities against a widely-used design approach that introduces a synthetic mismatch between the guide and its targets (Baseline; Methods). Top, distributions of the difference in predicted on-target and off-target activities (*A*(*p* | *T*_1_) − *A*(*p* | *T*_2_)) across the 100 target pairs. Bottom, break-down of the difference into predicted on-target activity (*A*(*p* | *T*_1_); bottom left) and predicted off-target activity (*A*(*p* | *T*_2_); bottom right). **(c)** Additional guide mismatches were introduced by the MEAs relative to the targets. Namely, plotted value is the percent of MEA-designed guides that have a mismatch against both *T*_1_ and *T*_2_ at positions around the SNP. (Positive positions are 3^*1*^ to the SNP; negative are 5^*1*^.) **(d)** Position of the SNP within the MEA-designed guides. For the majority of target pairs, the MEAs place the SNP within Cas13a’s mismatch-sensitive seed region. **(e)** Experimental results of guides designed with different approaches for identifying the S139N SNP in Zika virus and A437G antimalarial resistance SNP in *P. falciparum* at a target concentration of 10^8^ copies/μL. The title of each plot indicates the target the guides were designed to detect as the on-target. **(f)** Normalized fluorescence over time of guides designed to identify the K417N/T SNP in SARS-CoV-2 at 10^8^ copies/μL. **(g)** Normalized fluorescence over time of guides designed to identify each of the four dengue virus serotypes at 10^6^ copies/μL. **(h)** Normalized fluorescence over time of guides designed to identify SARS-CoV-2 lineages at 10^8^ copies/μL. For the Omicron BA.2 on-target, the WGAN-AM and baseline methods designed the same guide (purple). For the Ref on-target, the WGAN-AM and evolutionary methods designed the same guide (green). In **(f–h)**, there are multiple off-targets for each discrimination task, so the dotted off-target curve shows the maximum fluorescence of the guide across the off-targets computed at each time point (e.g., the off-target curves for K417 represent the fluorescence for the higher of the K417N and K417T targets at each time point in the reaction). **(i)** Evaluation of the effect of the tag-adjacent mismatch on rescuing guide activity when there are unfavorable (GN) nucleotides at the anti-tag region. Curves show the normalized fluorescence of guides detecting targets that have different nucleotide pairs at the anti-tag region, at 10^7^ copies/μL. All guides are identical to the targets at positions 1-27 in the protospacer. The colored lines represent guides with a terminal adjacent mismatch (at position 28), while the black dashed lines represent guides without a mismatch to the target. The dinucleotide above each plot indicates the first two nucleotides of that target’s anti-tag region, and the guide sequences represented in the schematic are reverse-complemented.

To computationally evaluate the performance of the MEAs in designing guides that identify SNPs, we considered a set of 100 pairs of randomly-generated on-target and off-target sequences, each differing by one nucleotide (Methods). We applied the MEAs to these target sets and also designed guides using the canonical strategy^24^ for this problem: this strategy places the SNP at position 26 of the guide and introduces a synthetic mismatch, at position 24 of the guide, to both the on-and off-target sequences (Methods). In silico, the MEA-designed guides exhibited similar predicted on-target activity to those designed by this synthetic mismatch method, but achieved lower off-target activity (Fig. 3b–d).

We benchmarked the MEAs’ variant identification performance experimentally, first by designing guides for six clinically-relevant SNPs and comparing them to the baseline canonical synthetic mismatch approach. The six SNPs include four associated with antimicrobial resistance in *Plasmodium falciparum* (Pfcrt K76T, Pfdhps A437G, Pfk13 C580Y, Pfmdr1 Y184F), one associated with microcephaly in Zika virus (PrM S139N), and one associated with immune evasion in SARS-CoV-2’s Spike protein (K417N/K417T)^26–32^. Experimentally, across nearly all of these tasks, the MEA-designed guides exhibited lower off-target activities while maintaining similar on-target activities to baseline guides (Fig. 3e–f and Supplementary Fig. 9a–c). Several baseline guides (including those for the S139, A437G, and K417T targets) had an off-target signal nearly identical to the on-target signal throughout the reaction, which complicates SNP identification; in contrast, the MEA-designed guides achieved an on-target signal 2–3 times above the off-target signal (Fig. 3e–f).

We further benchmarked the MEAs’ variant discrimination performance experimentally for designing guides that differentiate viral lineages, focusing on dengue virus (DENV) serotypes 1–4, and seven key SARS-CoV-2 lineages (Methods). In the DENV task, one serotype is the on-target set (e.g., DENV-1) and the other three serotypes comprise the off-target set (e.g., DENV-2, 3, 4). Both the MEA and baseline guides differentiated the DENV subtypes with high specificity at a target concentration of 10^8^ copies/μL, likely because there are DENV genomic regions with sufficient dissimilarity across serotypes (Supplementary Fig. 9d). However, when tested at a lower target concentration of 10^6^ copies/μL, the baseline guides exhibited low on-target fluorescence for serotypes 2 and 4, while the MEA-designed guides achieved, as desired, a high on-target fluorescence for those serotypes (Fig. 3g). In the SARS-CoV-2 task, we designed guides that differentiate the ancestral Wuhan lineage (i.e., Ref), Alpha, Beta, Delta, Gamma, Omicron/BA.1, and Omicron/BA.2 (Methods). The MEA-designed guides enabled robust lineage discrimination, while the baseline guides for Alpha, Gamma, and Ref had off-target activities that were nearly as high as their on-target activities throughout the reaction (Fig. 3h).

Overall, the WGAN-AM and evolutionary algorithms performed similarly in our two tasks, despite employing distinct techniques and creating different guides. On average, the evolutionary algorithm designed guides that are more distinct from observed sequences than the WGAN-AM’s guides (Figs. 2c, 3d, and Supplementary Fig. 5).

Next, we sought to understand if the MEAs leverage known rules governing Cas13a activity to enhance guide performance. Remarkably, we found that the MEAs implicitly learn Cas13a’s proto-spacer and mismatch preferences, without having them explicitly encoded, and rationally apply this biological understanding among our two design objectives to generate optimally-fit guides. For example, guide-target mismatches in close proximity to one another are known to decrease enzymatic activity^33–35^. When applied to the first objective of multi-target detection, the MEA-designed artificial guides reduced the number of adjacent guide-target mismatches at mismatch-sensitive positions relative to baseline guides, enabling superior sensitivity (Supplementary Fig. 10). In contrast, when applied to the variant identification objective, the MEAs introduced artificial mismatches within 4 nt of the SNP for the majority of target sets (WGAN-AM, 61%; evolutionary, 81%), reducing off-target activity (Fig. 3c). Additionally, for nearly 75% of simulated targets, the MEAs further optimized guide specificity by placing the SNP between positions 18 to 23, a region where mismatches are known^22^ to be highly deleterious to LwaCas13a activity (Fig. 3d).

Furthermore, we found that the MEAs take advantage of another biological property of Cas13a to design guides optimally suited for both objectives. Previous work has found that Cas13a guides exhibit reduced activity when targeting a genomic site with complementarity to Cas13a’s tag sequence, particularly targets with a `G` nucleotide at the position immediately 3^*′*^ to the protospacer (the protospacer-flanking site, or PFS)^35, 36^. In the variant identification objective, when the targeted SNP is a mutation from a non-`G` to a `G` nucleotide (as it is in the experimentally-tested S139N and C580Y targets), the MEAs positioned the `G` at the PFS to design guides with minimal off-target activity (schematized in Supplementary Fig. 11; experimental results in Fig. 3e and Supplementary Fig. 9a). However, in the multi-target detection objective, the MEAs exhibited an unexpected design decision. For genomic sites with a `G` at the PFS, the MEAs often introduced a mismatch directly adjacent to the tag sequence at position 28 in the protospacer (the same as position 1 in the spacer), which we term the tag-adjacent mismatch, or TAM. Intriguingly, among the genomic sites with a `G` at the PFS, the TAM was present in 45.7% of guides designed by the WGAN-AM algorithm and 76.2% of guides designed by the evolutionary algorithm (Supplementary Fig. 12).

We hypothesized that the TAM could represent a novel design strategy discovered by the MEAs that enables improved guide activity on targets with a `G` at the PFS, so we explored the TAM’s significance further. Mechanistically, the TAM may disrupt base pairing between the `G` at the PFS of the target and the `C` in the CRISPR RNA tag sequence, promoting the separation of the Cas13– target complex and thereby increasing collateral cleavage activity and the resulting fluorescent signal. To test our hypothesis, we designed a library of guides against targets having unfavorable nucleotides (`GN`) and favorable nucleotides (`AN`, `TN`, or `CN`) for the first two positions of the anti-tag region; the first nucleotide of the anti-tag region is the PFS (Methods; Fig. 3i). Guides with a TAM targeting a sequence with a `G` at the PFS exhibited substantially greater activity than guides without this TAM (Fig. 3i and Supplementary Fig. 13). Furthermore, the TAM conferred the greatest benefit when the targeted sequence was GT in the anti-tag region. This is mechanistically explainable: the first two nucleotides of Cas13a’s tag sequence complement `GT`, so the TAM’s disruption of this double base pairing may enhance activity more than for targets with a non-`GT` allele (Fig. 3i and Supplementary Fig. 13). As expected, introducing a TAM to guides targeting a sequence with a non-`G` at the PFS did not increase activity. To our knowledge, this design principle of introducing a mismatch in the spacer to rescue the activity of guides with extended tag:anti-tag complementarity has not been described before, suggesting that MEAs are capable of elucidating new design principles.

In this work, we demonstrate that MEAs can generate maximally-fit guide sequences for pathogen detection and surveillance, and can illuminate novel guide design rules. These algorithms leverage a predictive model to explore a sequence landscape, only evaluating ∼ 4,000 out of ∼ 10^16^ potential guides throughout their search process, yet are capable of designing artificial guides that outperform those derived directly from nature (Supplementary Note 1). Crucially, our approach is not trained on nor restricted to a specific design task, but rather can generate guides for any conditioned target set. The machine-learned models of guide activity that steer our optimization process enable our method to exploit features of Cas13a’s underlying biology (e.g., anti-tag and mismatch preferences) to generate guides that optimize diagnostic performance. During the early 2022 surge in COVID-19 cases, we applied a prototype version of our method to design guides that distinguish SARS-CoV-2 variants, demonstrating that MEA-designed panels achieve high concordance with next-generation sequencing in a clinical context^37^. We note that while the MEA-designed guides in our study achieved robust experimental performance, these methods rely upon predictive models of guide activity, which may return inaccurate predictions for guide-target pairs outside of the training domain. This is an area for further study. These methods can also reveal new, interpretable guide design rules for CRISPR-based technologies. The TAM proposed by the MEAs may enable Cas13a-based technologies to more efficiently target a broader range of sequences and improve our understanding of the enzyme’s nuclease activation mechanisms.

Our work suggests that MEAs can advance guide design for several technologies. In the realm of pathogen diagnostics, to our knowledge, all current design methods for nucleic acid technologies derive oligonucleotide sequences directly from pathogen genomes^38–41^. Artificial sequences stand to optimize the design of diagnostic sequences beyond CRISPR guide RNAs, including the primers and fluorescent probes employed in PCR, loop-mediated isothermal amplification (LAMP), and recombinase polymerase amplification (RPA). Beyond diagnostics, MEAs could design rationally mismatched guide RNAs for base editors to minimize bystander editing rates^23^. For certain CRISPR screens, where the objective is to introduce maximal sequence diversity into protein-coding genes, these methods could design guides that maximize the number of distinct edits made at the targeted sites^42^. Furthermore, while high-fidelity variants of CRISPR enzymes have lowered off-target editing rates^43^, MEAs could optimize guide RNA sequences to minimize these off-target effects further. Recent work has also highlighted the importance of accounting for human genetic diversity when designing CRISPR therapeutics^43, 44^, and MEAs are poised to generate guide sequences that optimally account for this diversity.

Generative methods are transforming the field of biological sequence design, with recent work focusing on generating long protein and DNA sequences^9, 11, 12^. Here, we show that generative design methods not only optimize shorter RNA sequences, such as CRISPR RNAs, but can simultaneously elucidate new mechanistic design rules and serve needs in pathogen surveillance.

## Acknowledgements

We thank Tien Nguyen, Jon Arizti Sanz, and Luca Pinello for valuable discussions and advice. This project was made possible by the Amazon Web Services Diagnostic Development Initiative and DARPA grant no. D18AC00006. H.C.M. was supported by NIH/NIAID grant no. K01AI163498 and USAID Prime grant no. 58-3022-2-031. S.M. and B.A.P. were supported by NIH/NIGMS grant no. T32GM144273. B.A.P. was also supported by NIH/NIGMS grant no. T32GM007753. C.M. was supported by INV-034761 from the Bill and Melinda Gates Foundation and the Princeton Catalysis Initiative. P.C.S was supported by HHMI, NIH/NIAID grant no. U19AI110818 and U01AI151812, CDC grant no. NU50CK000629 and 75D30122C15113, Flu Lab, and a cohort of generous donors through TED’s Audacious Project, including the ELMA Foundation, MacKenzie Scott, the Skoll Foundation, and Open Philanthropy.

## Author contributions

S.M. developed, implemented, and evaluated model-directed exploration algorithms for Cas13-based viral detection. P.P.P., N.L.W., C.M., and B.P. provided critical insights. H.C.M. initiated the project and performed some of the early methods development and analysis. S.M. and H.C.M. wrote the manuscript with input from other authors. H.C.M. and P.C.S. supervised the project.

## Methods

The model-directed exploration algorithms developed in this work pair (1) a predictive model of diagnostic guide activity with (2) exploration algorithms that search through a landscape of guide sequences. Together, they design guides that optimize a well-defined objective function.

A guide sequence is a ∼ 20 to 30 nucleotide (nt) nucleic acid sequence that binds to a target. In our applications, the guide is the 28 nt CRISPR RNA (crRNA) spacer sequence for use with CRISPR-Cas13a–based detection. It allows detection of a target by leading to a readout when the target is present.

### Predictive model of diagnostic guide activity

We previously developed the predictive models used in this work and they are described in detail in ref. 22. The input into these models is a 28-nt guide sequence and a 48-nt target sequence (having 10 nt of context on each site of the guide’s binding region). The output of the classification model *C*(*g* | *t*) ∈ [0, 1] can be interpreted as the probability that a given guide-target pair is active. The output of the regression model *R*(*g* | *t*) ∈ [− 4, 0] is the predicted activity of an active guide-target pair. Higher values of *C*(*g* | *t*) may suggest that a guide is more likely to bind to a given target, and higher values of *R*(*g* | *t*) may suggest that, after binding, a guide’s signal on a given target will grow faster.

In this work, we combined the classifier and regression model’s predictions to define a weighted activity metric, *A*, which follows from the law of total probability:

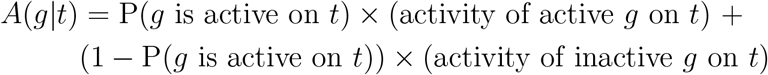

In our case, *C*(*g* | *t*) = P(*g* is active on *t*), *R*(*g* | *t*) = activity of active *g* on *t*, and − 4 = activity of inactive *g* on *t* because guides that are inactive have a near-zero signal, which is represented in our dataset by an activity of *−*4, the lower bound of measured signal.

Thus, for a given guide-target pair, we computed *A*(*g*|*t*) as follows:

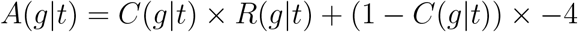

### Objective functions for diagnostic and surveillance applications

Model-directed exploration algorithms can design a guide sequence that optimizes a well-defined objective function across a genome’s sequence variation.

We focus on designing guides that optimize two specific objective functions. These objectives have applications in diagnostic testing and pathogen genomic surveillance.

### Multi-target detection

In the first application (“multi-target detection”), our goal is to design guides that are maximally active across genome sequence diversity. As such, the objective function for this application is:

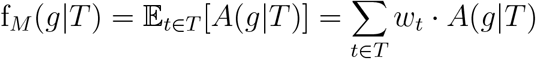

where *T* is the set of genome sequence targets and *t ∈ T* is a single target.

Genome sequence diversity varies spatially and temporally, so the *w*_*t*_ represents a prior probability of encountering a target *t* ∈ *T* . We set a uniform prior distribution (assuming that all *w*_*t*_ are equal 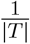) over the target genome sequences, following the choice in ref. 22, but one could modify *w*_*t*_ as certain lineages of a pathogen become more or less prevalent in a geographic region of interest or over time.

Additionally, one could also consider a defining more complex objective functions (such as the 10^th^ percentile of the distribution of activity across genomic diversity), although these functions may be non-differentiable and more difficult for certain exploration algorithms to maximize.

### Variant identification

In the second application (“variant identification”), our goal is to design a diagnostic guide *g* that can optimally differentiate between closely related sequences, maximizing activity against one set of targeted sequences (on-target) while minimizing activity against a separate but homologous set of non-targeted sequences (off-target). In our notation, *T*_1_ represents the on-target sequences and *T*_2_ represents the off-target sequences.

Initially, we considered a simple objective function: the difference between the expected on-target activity and expected off-target activity, 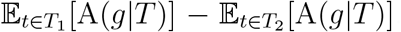. Although many guides designed using this objective had a large difference between the expected on-target activity and expected off-target activity, they were often inactive or only marginally active on the on-target sequences—making these guide designs unusable—because the function does not enforce that the guide’s activity on the on-target set of sequences be sufficiently high for detection.

Thus, we defined an objective function for the variant identification application that is maximized only when the designed guide has high activity on the on-target sequences and low activity on the off-target sequences. We based this objective function upon the logistic function so it is differentiable. The objective is defined below:

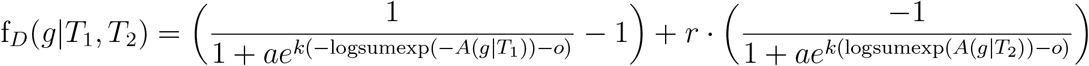

where *o, a, k*, and *r* are all parameters that modulate the slope and curvature of the objective function. The values of these parameters were determined through a random search procedure described in the hyperparameter search section of the Methods. We use logsumexp(*A*(*g* | *T*_*i*_)) as shorthand for logsumexp({*A*(*g* | *t*) ∀ *t* ∈ *T*_*i*_ }), where logsumexp(*x*_1_, …, *x*_*n*_) = log(exp(*x*_1_) + + exp(*x*_*n*_)) is a smooth approximation to the maximum.

The value of f_*D*_(*g* | *T*_1_, *T*_2_) increases as the minimum activity of the guide across the on-target set increases and the maximum activity of the guide across the off-target set decreases. A visualization of this objective function is included in Supplementary Fig. 14.

### Exploration algorithms

The above objective functions define the fitness of a guide for two types of detection tasks. Our goal is to design maximally-fit guide sequences—those that maximize the value of these objective functions. We developed two algorithms, a Wasserstein generative adversarial network–activation maximization (WGAN-AM) approach and an evolutionary algorithm, that search over a landscape of guide sequences to generate these maximally-fit guides.

### Conditional generative adversarial network with activation maximization (WGAN-AM)

A generative adversarial network (GAN) is an unsupervised deep learning framework that approximates an underlying data-generating distribution^45^. A well-trained GAN can generate new samples of data (e.g., photographs or genomic sequences) that have properties similar to those of the training set.

We use a Wasserstein GAN (WGAN) inspired by the application in ref. 25, but with one key difference. Our WGAN uses a conditional GAN, in particular conditional on a given target set^46^. The generator network, G(*z* | *T*), generates a guide sequence for the given target set. This difference allows our method to generalize well across different target sets, which contrasts to using a GAN that has been fit to data derived from a single target set. The WGAN uses a latent variable *z* as an input, and *z* can be modulated to explore the sequence space and generate different guide sequences for a given target set.

The generator network of the WGAN proceeds in five stages. First, the input latent vector *z* is upsampled to a vector of | *z* | × guide length. In our implementation, the latent vector is 10-dimensional and the guide has a length of 28, so the resulting upsampled vector has length 280. Second, this vector is reshaped into a matrix with dimensions guide length × latent channels (28 by 10). Third, this matrix is padded with a 10 by 10 matrix of zeros on each side such that the dimensions of the resulting padded matrix is 48 by 10. Fourth, the 48 by 10 padded matrix is appended to the one-hot-encoded consensus of a 48-nt region of the target, which has dimensions of 48 by 4. The resulting matrix has dimensions 48 by 14. This makes the generator conditional a given set of targets, because a representative sequence (the consensus) is directly concatenated in this operation. Fifth, this matrix is passed through three residual blocks and 4 convolutional filters with stride 1 and width 2 to create a 48 by 4 matrix. Finally, a softmax is applied such that the resulting matrix represents the probability of having a base at each position, and the 10 by 4 context is removed from either side. The resulting 28 by 4 matrix represents the 28 nucleotide guide sequence, which is the guide that we sought to design. Supplementary Fig. 15a schematizes the architecture of the generative network of our WGAN.

We trained the WGAN on the same set of active guide-target pairs that our previously-developed CNN regression model was trained on^22^. These guide-target pairs were experimentally demonstrated to be active via the CARMEN-Cas13 system^22^. Thus, the generator outputs guides with measurable activity against the conditioned target. We trained the WGAN via the WGAN-gradient penalty method in ref. 47. We used a batch size of 32 and trained the critic 5 times for each batch on which the generator was trained. We used the following hyperparameters for the Adam optimizer^48^: *α* = 0.0001, *β*_1_ = 0.9, *β*_2_ = 0.999, and ϵ = 10^*−*7^.

The generator network G(*z* | *T*) captures the high-level features that make a guide active against a given target set, and is capable of generating an artificial guide sequence given a target set. To generate maximally fit diagnostic guides, we employed a joint activation maximization framework^25^.

In this joint framework, the WGAN’s generator, G(*z*|*T*), transforms the latent vector *z* into a candidate guide sequence *g*, and the objective function, f(*g*|*T*), computes the guide’s fitness.

To generate optimal guides, we compute the gradient of the guide’s fitness with respect to the latent vector *z*, and use the Adam optimizer^48^ to take steps in the direction of this gradient:

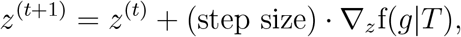

where 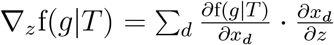 and *d* is one of the 10 dimensions of the latent vector.

This procedure allows us to perform a continuous search over the latent space to generate guide sequences that have optimal fitness. To more thoroughly explore the fitness landscape and account for the fact that several local minima may exist in the fitness landscape, we perform multiple searches over the latent space with each search starting at a different random starting point (sampled from a normal distribution) in the latent space. A pseudocode overview of the WGAN-AM algorithm is available in Supplementary Fig. 1 and a high-level schematic is presented in Fig. 1c.

The WGAN-AM algorithm’s generated guide sequences are implicitly constrained. Since the WGAN’s generator network approximates the conditional probability distribution P(*g* | *T*) of the guide-target pairs in the training set, the guide sequences output by the WGAN’s generator network have similar properties (e.g., mismatch positions and mismatch types) to the guide sequences used to train the regression model. Thus, the WGAN-AM algorithm searches regions of the fitness landscape in which the regression model is likely to accurately predict guide activity. Because of the generator network’s structure, it is unlikely that the WGAN-AM algorithm will design guides that are so different than the guide sequences in the training set that the regression model’s predictions are no longer accurate and the model behaves pathologically. For example, we would not expect the WGAN-AM algorithm to generate guides that have a much higher number of mismatches against their target sets than those in the training data for the predictive models.

### Evolutionary algorithm

We also tested an unconstrained search algorithm that could reach any region of the fitness landscape— an evolutionary algorithm. Evolutionary algorithms are biologically-inspired optimization algorithms that are widely used in computer science to search across a domain of candidate solutions.

They have been applied to design biological sequences that exhibit a desired property, such as peptide sequences that have maximal antimicrobial activity^2, 49–51^.

We build upon this work to develop an evolutionary algorithm that searches over our fitness landscape of guide sequences by performing iterative rounds of fitness evaluation, selection, and mutation^51^.

First, we initialize a population of guide sequences *G*_0_ by extracting all of the guide-length sequences at the genomic site of interest. Then, we use the objective function f(*g* | *T*) to evaluate the fitness of each of these candidate guides. We sample *S* parent guides from this population with a probability 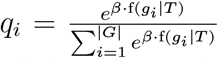, where *β* represents the selection intensity^51^. We then mutate the randomly-sampled guides with a mutation frequency *γ*. Specifically, for each position in the guide, we randomly sample a value from the standard uniform distribution, *x*_*i*_ ∼ Uniform(0,1). If *x*_*i*_ *< γ*, we mutate the nucleotide at that position (e.g., ‘A’) to one of the other nucleotides (e.g., ‘C’, ‘G’, or ‘U’) with uniform probability. Finally, we add the resulting child guides to the guide population. We repeat these rounds of fitness evaluation, selection, and mutation for several generations until the limit on the number of calls to the objective function, *L*, is exceeded. We return the guide sequences in the final population.

Throughout the search process, the MEAs are capable of efficiently recognizing the positions at which fitness increases if the guide diverges from the consensus sequence, but they don’t explicitly evaluate each possible nucleotide at those positions. Thus, after the last generation (of the evolutionary algorithm) and after the last outer round (of the WGAN-AM algorithm), the algorithms perform a local search, where they identify positions at which the generated guide differs from the consensus, mutate the guide to the other possible nucleotides at each of these positions, and then select the resulting guide with the best fitness.

A pseudocode overview of the evolutionary algorithm is available in Supplementary Fig. 2, and a high-level schematic is presented in Fig. 1d.

We implemented and tested all models using TensorFlow 2.8.0^52^ and FLEXS 0.2.8^2^ in Python 3.7.10.

### Hyperparameter Search

We implemented a random search procedure to determine the optimal hyperparameters for our design algorithms. Because we developed two model-directed exploration algorithms (WGAN-AM and evolutionary) and we have two objective functions (f_*M*_ (*g* | *T*) and f_*D*_(*g* | *T*_1_, *T*_2_)), four sets of hyperparameters were determined by random search.

#### Multi-target detection objective function

For the multi-target detection objective function, f_*M*_ (*g* | *T*), we constructed a random search dataset of genomes from NCBI’s GenBank database^53^. We randomly sampled ten viral families from a list of all viral families that have at least one vertebrate-infecting species. Then, within each of these ten families, we randomly sampled one vertebrate-infecting species that has at least 100 complete genomes. The ten selected viral species were Primate T-lymphotropic virus 1 (NCBI taxonomic identifier (taxid): 194440), avian orthoreovirus (taxid: 38170), alphapapillomavirus 10 (taxid: 333754), yellow fever virus (taxid: 11089), eastern equine encephalitis virus (taxid: 11021), pigeon circovirus (taxid: 1414603), SFTS phlebovirus (taxid: 1933190), human respovirus 3 (taxid: 11216), human metapneumovirus (taxid: 162145), and Betacoronavirus 1 (taxid: 694003). For each of these ten viral species, we used ADAPT^22^ to compile and curate complete genomes and used MAFFT^54^ create an alignment of the curated genomes. We randomly selected ten guide-length sites from each of the ten alignments, and extracted all of the sequences at each genomic site to create a target set *T* . In summary, we constructed 100 target sets from 100 genomic sites across ten different viral species—allowing our hyperparameter search process to reflect viral diversity.

We created 100 sets of hyperparameters for the WGAN-AM algorithm and the evolutionary algorithm by randomly sampling from the distributions listed below. For each set of hyperparameters, we ran the WGAN-AM and evolutionary algorithms across all 100 target sets, and computed the mean fitness of the guides across the target sets 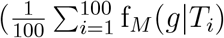, for a set of targets *T*_*i*_ at each site). We ultimately selected the set of hyperparameters that the maximized mean guide fitness.

#### Variant identification objective function

For the variant identification objective function, f_*D*_(*g* | *T*_1_, *T*_2_), we created a synthetic random search dataset. We randomly generated 50 DNA sequences of length 150 to form the on-target sets 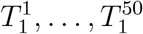. For each of these DNA sequences, we mutated the base in the center of the sequence (position 75) to another randomly-selected base. These mutated DNA sequences comprised the off-target sets, 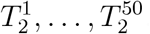. These pairs of on-target and off-target sets 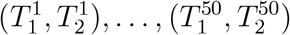 represent two different sequences, one with a single nucleotide mutation and one without a single nucleotide mutation.

We created 100 sets of hyperparameters for the WGAN-AM algorithm and the evolutionary algorithm by randomly sampling from the distributions listed below. Furthermore, since the variant identification objective function f_*D*_(*g* | *T*_1_, *T*_2_) has hyperparameters as well, we created 100 sets of objective function hyperparameters by randomly sampling from the distributions also listed below.

For each set of hyperparameters, we ran the WGAN-AM and evolutionary algorithms across the 50 pairs of on-target and off-target sets 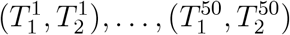 in a sliding window fashion so that we would examine a guide placing the SNP at each position of the guide (window length = 28 nt, window stride = 1 nt; selecting the optimal of the 28 guides) to design 50 diagnostic guides. Because the objective function f_*D*_(*g* | *T*_1_, *T*_2_) has its own hyperparameters *k, o, a*, and *r*, it would not be appropriate to directly compare the fitness of the generated guides across the different hyperparameter sets. Since our goal is to design guides that have high on-target activity and low off-target activity, we selected the set of hyperparameters that maximized the mean difference between predicted on-target and off-target activity 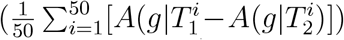 while simultaneously generating guides with a high predicted on-target activity 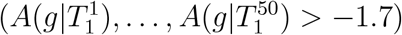.

#### List of hyperparameters and search intervals

WGAN-AM algorithm parameters determined through random search:

- Number of starting points, *r*_*outer*_ *∼* uniform in [5, 35] for multi-target detection objective, *r*_*outer*_ *∼* uniform in [5, 15] for variant identification objective
- Number of steps in each search of the latent space, *r*_*inner*_ *∼* uniform in [50, 325] for multi-target detection objective, *r*_*inner*_ *∼* uniform in [50, 230] for variant identification objective
- optimizer learning rate, *α ∼* log-uniform in [10^*−*2^, 10^2^]

Evolutionary algorithm parameters determined through random search:

- Selection intensity, *β* ∼ log-uniform in [10^*−*1.2^, 10^2.2^] for multi-target detection objective, *β* ∼ log-uniform in [10^*−*2^, 10^2^] for variant identification objective
- Mutation rate, *γ ∼* uniform in 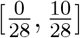 for multi-target detection objective, *γ ∼* uniform in 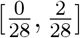 for variant identification objective
- Fraction of children in new generation, *j ∼* uniform in [0.1, 0.9]
- Population size, *S* ∼ uniform in [50, 300] for multi-target detection objective, *S* ∼ uniform in [50, 250] for variant identification objective
- Limit on number of calls to the objective function, *L* = 1500 to control the runtime of the algorithm

Variant identification objective function parameters determined through random search:

- *a ∼* uniform in [0.1, 9.1]
- *k ∼* uniform in [*−*4, *−*1]
- *r ∼* uniform in [0.1, 10.1]
- *o ∼* uniform in [*−*4.5, *−*1]

#### Random search-determined hyperparameter values

The hyperparameters determined through these random search procedures are as follows:

##### Multi-target detection

WGAN-AM algorithm

- *r*_*outer*_ = 28
- *r*_*inner*_ = 275
- *α* = 1.540127

Evolutionary algorithm

- *β* = 0.077373
- *γ* = 0.003362
- *j* = 0.794996

##### Variant identification

WGAN-AM algorithm

- *r*_*outer*_ = 8
- *r*_*inner*_ = 144
- *α* = 0.632998
- *a* = 3.769183

Evolutionary algorithm

- *β* = 2.201796
- *γ* = 0.029049
- *j* = 0.893401
- *S* = 119
- *S* = 87
- *L* = 1500
- *k* = *−*3.833902
- *r* = 2.973052
- *o* = *−*2.134395
- *a* = 5.897292
- *k* = *−*2.857755
- *r* = 1.736507
- *o* = *−*2.510856

### Computational benchmarking of model-directed exploration algorithms

To characterize the ability of the WGAN’s generator network to introduce mismatches at different positions in the guide and between different alleles, we ran a simulation. We computed the consensus sequence at each of the 100 genomic sites used in the multi-target detection random search. For each consensus sequence *C*_1_, …, *C*_100_, we randomly sampled 500 latent variables *z* ∼ *N* (0, 1) and generated 500 guides conditioned on that consensus sequence using the generator network G(*z* | *C*_*i*_). We computed the Euclidean norm of the latent variable as well as the the Hamming distance between the WGAN-AM generated guide and the consensus sequence, which is the number of positions at which the two guides have different nucleotides.

We found that that as the Euclidean norm of the latent variable *z* increases, the guide generated by the WGAN’s generator network has a higher Hamming distance from the sequence that it was conditioned upon. In other words, as the latent variable *z* gets farther from the origin, the generator introduces more mutations into the generated guide. This property is illustrated in Supplementary Fig. 15b. Thus, the latent space *z* has an interpretable biological meaning.

Additionally, to benchmark the algorithms’ ability to design guides for the variant identification objective, we created a synthetic benchmarking dataset. In order to avoid biasing the sequence composition towards a limited set of pathogen genomes and evaluate the performance of the methods across a large distribution of sequences, we randomly generated 100 pairs of DNA sequences, each pair consisting of targets that differ by one nucleotide. The MEAs were run on this dataset and the results are shown in Fig. 3b–d.

### Designing diagnostic guides for experiments

We applied our model-directed exploration algorithms to design diagnostic guides for both the multi-target detection and variant identification applications.

### Multi-target detection experiments

For the multi-target detection application, we selected five RNA viruses that are of public health interest and cause significant morbidity and mortality globally. We used ADAPT^22^ to download and curate complete genomes for dengue virus (taxid: 12637), influenza A virus (segment 2, taxid: 138949), and enterovirus B (taxid: 138949) on August 30, 2021 from NCBI’s GenBank^53^. For SARS-CoV-2, we downloaded all complete genomes available on GISAID^55^ on June 28, 2021, removed the sequences flagged as ‘low quality’, and randomly sampled 10,000 of the genomes. For Lassa virus (segment S, taxid: 11620), we directly downloaded complete genomes from GenBank^53^ using the following search criteria: ‘txid11620[Organism:exp] AND “segment S” AND “complete sequence”’ on August 31, 2021. We used MAFFT to generate alignments for these five viral species^54^.

The predictive models (*R*(*g* | *T*) and *C*(*g* | *T*)) employed in this work require an input of a 48 nt target sequence (a 28 nt guide-binding target sequence plus 10 nt of context on each side). Thus, we extracted 48 nt sliding windows along the genome and considered the target set *T* at that genomic site. We removed the genomic sites with less than 10 valid sequences (sequences that are a contiguous 48 nt and have no ambiguity or gaps). Both of the model-based exploration algorithms were run on all of these target sets using the multi-target objective function, f_*M*_ (*g* | *T*), to design guides for these genomic sites.

To benchmark the performance of our model-based exploration algorithms against current methods, we used ADAPT to design guides at each of these genomic sites. ADAPT is software suite that employs machine learning and combinatorial optimization to design diagnostic guides and is a state-of-the-art in the field^22^. When running ADAPT, we used the arguments ‘`-w 28 -gl 28 --predict-cas13a-activity-model --obj maximize-activity -hgc 1 --cluster-threshold 0.3`’ with`SEARCH-TYPE` set to`sliding-window`, such that the CNN-based predictive model was used and ADAPT was restricted to designing one guide at each site. We refer to this usage of ADAPT as “model-based choice” (MBC), since the predictive model is being used to select a maximally fit sequence present in the ground set.

Furthermore, we computed the consensus of the sequence alignment at each of these genomic sites to represent a simple baseline approach to guide sequence design. These guides are called “consensus guides” throughout the text.

We sought to appropriately compare the performance of the guides designed by the baseline methods (MBC and the consensus) with the performance of the guides designed by our model-directed exploration algorithms (WGAN-AM and evolutionary). It is not tractable to experimentally test the tens of thousands of diagnostic guides designed. Thus, for every genomic site, we computed a relative performance metric, RP_*i*_, the summed fitness of the guides designed by the model-directed exploration algorithms subtracted by the summed fitness of the guides designed by the baseline methods.

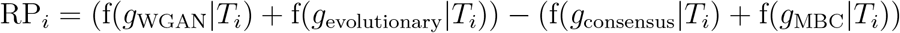

Previous studies^22,24^ have shown that LwaCas13a exhibits reduced activity when targeting a genomic sites with a`G` at the PFS, so we only considered genomic sites with a non-`G` at the PFS for experimental testing. For each viral species, all genomic sites with a non-`G` at the PFS were sorted into quantiles by their respective RP_*i*_, so genomic sites with the highest RP_*i*_ were in the fourth quartile. We randomly sampled two sites from the fourth quartile, and randomly sampled one site from the first quartile for each viral species. This method for selecting genomic sites allows us to experimentally test sites at which the guides designed by the model-directed exploration algorithms were likely to have superior performance than the guides designed by previous methods, as well as sites at which guides designed by previous methods were likely to have similar or better performance than the guides designed by the model-directed exploration algorithms. Each panel with experimental multi-target detection results shows one such site.

To computationally characterize guide performance, we computed the coverage of the guides designed by the MEAs and baseline methods across different genomic windows in the viral pathogens, as shown in Fig. 2a and Supplementary Fig. 3.

A guide *g* was considered to detect a target if it was predicted to be active by the classification model (*C*(*g* | *t*) *>* 0.577467) and was predicted to be in the top quartile of activity by the regression model (*R*(*g* | *t*) *>* − 1.2801363), based on the “highly active” criterion used in ref. 22. The coverage was computed by determining the percentage of targets a guide detects in the given target set.

We computed the predicted guide activity across all targetable genomic sites in these pathogens, as shown in Fig. 2b and Supplementary Fig. 4. Targetable genomic sites are defined as sites at which the baseline methods had meaningful activity 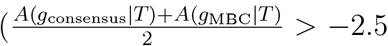; in practice, it would not make sense to target sites that fall below this minimal threshold). In these figures and in all boxplots throughout the manuscript, the box represents the first and third quartiles (*Q*1 and *Q*3) and the whiskers extend to from *Q*1 *−* 1.5 *· IQR* to *Q*3 + 1.5 *· IQR*.

### Variant identification experiments

For the variant identification application, we sought to design diagnostic guides that could differentiate between dengue virus (DENV) serotypes 1–4 and the SARS-CoV-2 WHO-designated variants of concern. For DENV, we downloaded all available complete genomes for serotypes 1-4 from the Virus Pathogen Database and Analysis Resource (ViPR)^56^ on February 1, 2022. We used MAFFT^54^ to generate an alignment of the genomes.

For SARS-CoV-2, we downloaded the ‘full length variant alignment’ from GISAID on January 22, 2022. This alignment contains representative sequences for the targeted SARS-CoV-2 lineages: Wuhan reference, Alpha (B.1.1.7), Beta (B.1.351), Gamma (P.1), Delta (B.1.617.2), Omicron (BA.1), and Omicron (BA.2). The GISAID accessions for these reference genomes are `EPI ISL 402124`, `EPI ISL 674612`, `EPI ISL 940877`, `EPI ISL 2777382`, `EPI ISL 1758376`, `EPI ISL 6914029`, and `EPI ISL 7190366`, respectively. We referenced analyses on the CoVariants^57^ and outbreak.info^58^ websites to confirm that the insertions, deletions, and mutations targeted by our guides were highly conserved and were present in at least 85% of genomic sequences of each variant.

The model-directed exploration algorithms were run on the DENV and SARS-CoV-2 alignments described above to design guides that were specific to each serotype/genotype/lineage.

To benchmark the performance of the model-directed exploration algorithms for variant identification of lineages (beyond a single SNP), we implemented a baseline approach. This baseline approach designs guide sequences by finding the 28-nt subsequence of the on-target consensus that has the highest Hamming distance from the off-target consensus and has an average nucleotide identity of at least 90% across the on-target genomes. This approach mimics the strategy (usually performed by hand) of identifying regions that are conserved among the on-target genomes but divergent from the off-targets.

For the antimalarial resistance panel, we downloaded reference sequences for the *P. falciparum* genes *Pfmdr1, Pfcrt, Pfk13, Pfdhps* from GenBank accessions `HQ215532.1`, `LC498250.1`, `KT328114.1`, and `KE123491.1`, respectively. For the K417N/T identification task, we used the same SARS-CoV-2 sequences that we previously downloaded from GISAID, as described above. For the Zika virus S139N identification task, we downloaded the reference sequence from GenBank accession `MT483911.1` . We extracted the 250 nt genomic regions encapsulating the sites of interest in each gene and used them as the target sequences.

The model-directed exploration algorithms were run to design guides for each of the 4 single nucleotide polymorphisms (SNPs) of interest in the *P. falciparum* genome, the K417N/T SNP in SARS-CoV-2, and the S139N SNP in Zika virus. We also sought to benchmark our algorithms against a baseline. The “synthetic mismatch” strategy (where the SNP is placed at position 26 of the protospacer and a mismatch at position 24 is introduced against both the derived and ancestral targets) is currently the standard approach for designing Cas13a guides that identify SNPs^24, 59^. The synthetic mismatch creates a double mismatch against the off-target sequence (positions 24 and 26), but only one mismatch against the on-target (position 24). We employed the synthetic mismatch strategy to design guides that target these 4 SNPs of interest, enabling us to benchmark the performance of our methods against the current standard.

### Tag-adjacent mismatch experiments

As discussed in the main text, the model-directed exploration algorithms often introduce a mismatch at position 28 in the guide when targeting a genomic site with `G` nucleotide at the protospacer-flanking site (Fig. 3). We refer to this mismatch the “tag-adjacent mismatch” (TAM) and hypothesized that it may enhance guide-target activity.

To determine if this was the case, we designed an experimental library consisting of targets that were representative of viral sequence diversity. We randomly sampled four viral families from a list of all viral families that have at least one vertebrate-infecting species. Then, within each of these four families, we randomly sampled one vertebrate-infecting species that has at least 100 complete genomes. The four selected viral species were primate tick-borne encephalitis virus (NCBI taxid: 11084), human mastadenovirus D (taxid: 130310), hepatitis B virus (taxid: 10407), and Zaire ebolavirus (taxid: 186538). For each of these viral species, we downloaded all available complete genomes from GenBank^53^ on February 15, 2022 and used MAFFT^54^ to create an alignment. We randomly selected a genomic site with a `G` nucleotide at the PFS in each of the four alignments and used this as our target sequence. We mutated the `G` nucleotide at the PFS in the original target sequence to create three additional targets with a non-`G` nucleotide at the PFS. Furthermore, since the nucleotide directly 3^*′*^of a site with a `G` at the PFS also impacts activity^22^, we additionally mutated the original target sequence to create targets that have an anti-tag region of `GC`, `GG`, `GA`, and `GT` .

For each of the four species, we designed the guide sequence without a TAM by simply extracting the 28 nt protospacer-binding region from the original target sequence. We also designed three guide sequences with the TAM by mutating the nucleotide at position 28 of the guide’s protospacer to the other three possible bases at that position.

#### Preparing target sequences for experiments

To experimentally test the performance of our guides, we first designed target sequences against which we would test guides. We designed these targets to be representative of viral genomic diversity. We ran ADAPT’s `pick test targets.py` program developed in ref. 22, with the following arguments: `--num-representative-targets 5 --min-target-len 250` . This script extracts a region of the alignment that encapsulates the guide-binding sites and is at least 250 nt long. Then, it clusters the resulting sequences and determines the medoid of each cluster, which are used as the representative target sequences. At all of the genomic sites we experimentally tested, the targets represented at least 95% of the total genomic diversity.

We added a T7 promoter sequence (5’-GAAATTAATACGACTCACTATAGGG-3’) followed by a positive control sequence (5’-CACTATAGGGGCTCTAGCGACTTCTTTAAATAGTGGCTTAAAATAAC-3’) to the 5’ end of every target sequence. In each experiment, we included a crRNA with a protospacer that was complementary to this positive control sequence (5’-GCTCTAGCGACTTCTTTAAATAGTGGCT-3’). This positive control enabled us to verify that the target sequences were synthesized correctly. *P. falciparum*’s genome may pose DNA synthesis challenges because it is `GC` -poor, so we added a second positive control sequence (5’-GAATGGAAGCACCGAGAGTATATGAAGATCTTCATGTGTGCAAAAGAATGGTAAAGCAGAGAAGGAGC-3’) to the 3^*′*^end of all the *P. falciparum* target sequences. In each experiment that had *P. falciparum* targets, we included crRNA with a protospacer that was complementary to this positive control (5’-TTCTTTTGCACACATGAAGATCTTCATAT-3’).

#### Preparing guide sequences for experiments

All of the methods developed and used in this work (including the model-based exploration algorithms and the baseline methods) output guide sequences in the frame of Cas13’s protospacer. To prepare the guide sequences for experiments, we took the reverse complement of them to transform them to the frame of the CRISPR-Cas13 spacer and added the LwaCas13 direct repeat (5’-GAUUUAGACUACCCCAAAAACGAAGGGGACUAAAAC-3’) to the 5^*′*^end of the crRNA sequence.

### Experimentally evaluating diagnostic guide designs

#### Experimental methods

We used the mCARMEN^37^ platform to experimentally test the performance of diagnostic guides. Briefly, mCARMEN is a CRISPR-based diagnostic technology that uses a Fluidigm microfluidic chip to enable highly multiplexed testing of dozens of diagnostic guides against dozens of RNA targets. We followed the protocol described in the “General mCARMEN procedures” section of ref. 37, with the following modifications:

All DNA targets were ordered as gBlocks from Integrated DNA Technologies. All crRNAs and the quenched synthetic fluorescent RNA reporter (`FAM/rUrUrUrUrUrUrU/3IABkFQ/`) were also ordered from Integrated DNA Technologies.

The DNA targets were serially diluted to the concentrations of 10^10^, 10^9^, and 10^8^ copies/μL, and 1.43μL of each target served as input to each sample mix. The quenched synthetic fluorescent reporter was included in the sample mix at a concentration of 500 nM. The crRNAs were included in the assay mixes at a concentration of 212.5 nM, and LwaCas13 from GenScript was included in the assay mix at a concentration of 42.5 nM.

After chip loading, the Fluidigm Biomark HD was set to a constant temperature of 37 °C and was used to image the IFC on the FAM and ROX channels every five minutes for three hours.

#### Analysis of experimental data

In order to robustly characterize the performance of each of the guide-target pairs, we computed reference-normalized background-subtracted fluorescence values, as was done in refs. 37 and 22.

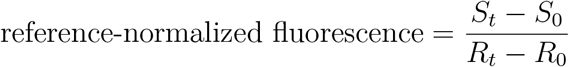

where *S*_*t*_ is guide-target pair’s FAM signal at time point *t, S*_0_ is the guide-target pair’s FAM signal at time point 0, *R*_*t*_ is guide-target pair’s reference ROX signal at time point *t*, and *R*_0_ is the guide-target pair’s reference ROX signal at time point 0.

To compute the reference-normalized background-subtracted fluorescence of each guide-target pair at time *t*, we subtracted the reference-normalized fluorescence of the no-template control at time *t* from the reference-normalized fluorescence of the guide-target pair at time *t*. Thus, if a guide-target pair has a reference-normalized background-subtracted fluorescence greater than 0, it has a fluorescence that is higher than the no-template control signal.

In the heatmaps of fluorescence, we plotted the reference-normalized background-subtracted fluorescence at the one hour timepoint. In the kinetic curves of fluorescence, we plotted all the reference-normalized background-subtracted fluorescence values collected throughout the course of the reaction (*t* = 0 to 180 minutes).

## Code availability

The MEA algorithms are available under the MIT license at https://github.com/broadinstitute/mea-cas13. The repository contains detailed instructions on how to run the MEAs on any input genomic sequences.

The code used to run the analyses in this manuscript is available at https://github.com/broadinstitute/mea-cas13-analysis.

## Supplementary Note 1

This note estimates the size of the sequence landscape that must be searched over to generate artificial guides.

The total number of possible 28-nt guides is 4^28^ *≈* 7 *×* 10^16^.

Optimally fit guide sequences would be nearby in sequence space to the complement of their target sequence. If *M* is the upper limit on the number of mismatches an optimal guide can have to this sequence, there are 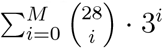 in this target-adjacent guide set. This totals to *∼*2.5 *×* 10^7^ and ∼ 2.9 × 10^9^ for *M* = 5 and *M* = 7, respectively. In many applications, we would explore this space not just once, but at every site in a targeted genome (tens to hundreds of thousands of sites for a typical viral genome); moreover, if a set of targets has high variation, we would explore the space around many observed alleles of the target. It would be inefficient to explore this vast space of potential guide sequences with an exhaustive, brute-force search.

## Supplementary Figures

**Supplementary Figure 1.**
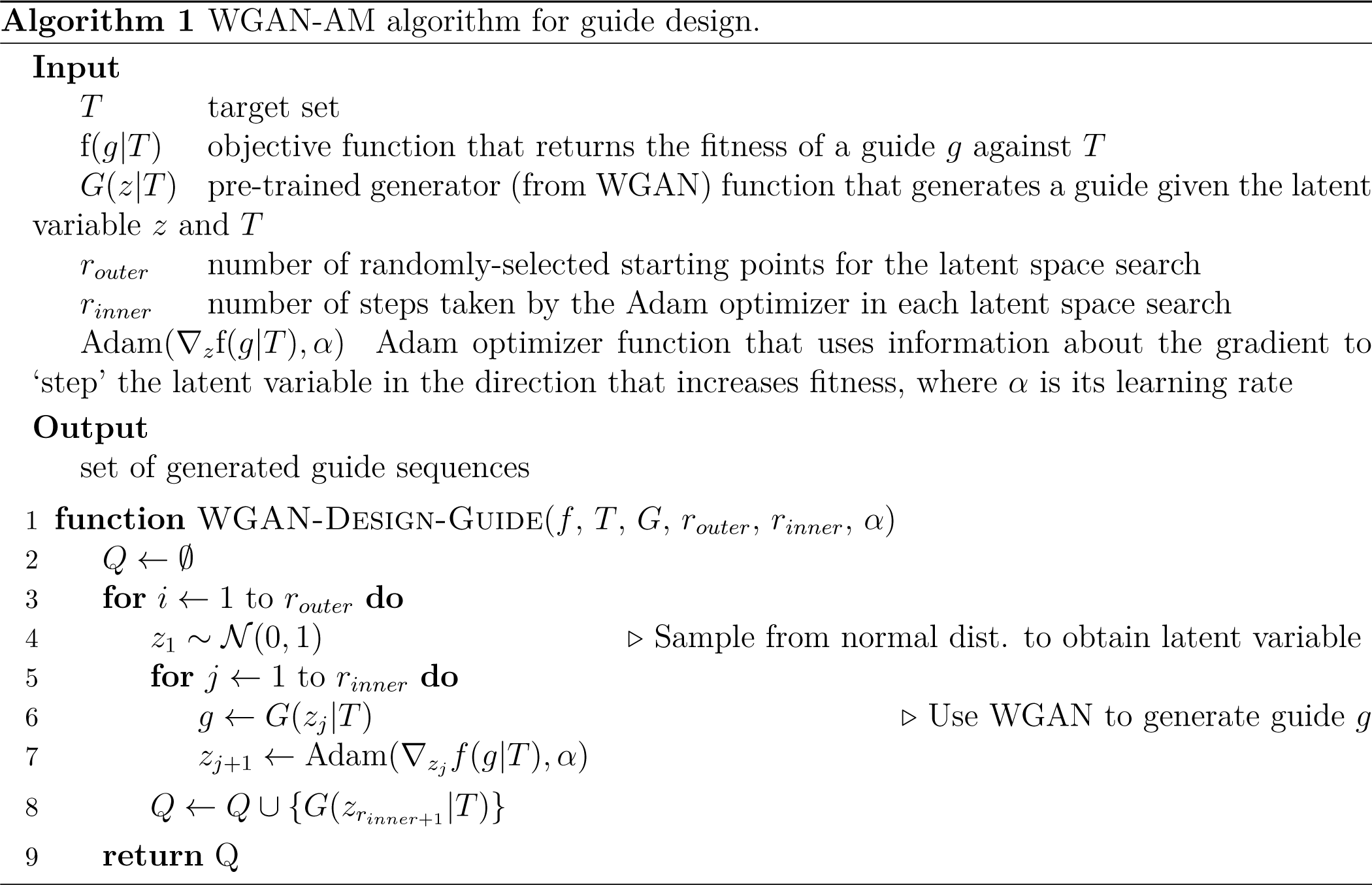
Pseudocode of the WGAN-AM algorithm. The WGAN-AM algorithm is further described in the Methods section.

**Supplementary Figure 2.**
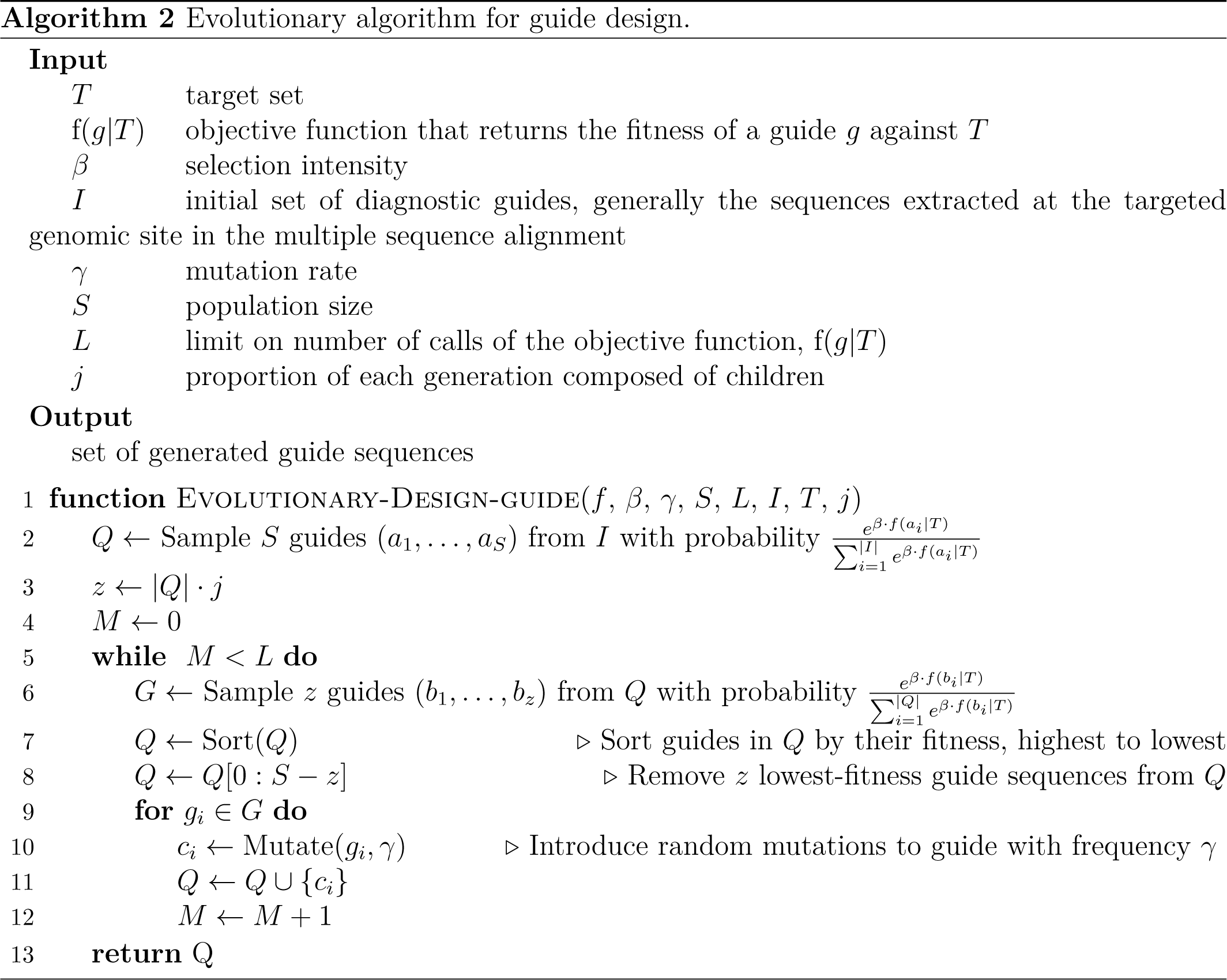
Pseudocode of the evolutionary algorithm. The evolutionary algorithm is further described in the Methods section.

**Supplementary Figure 3.**
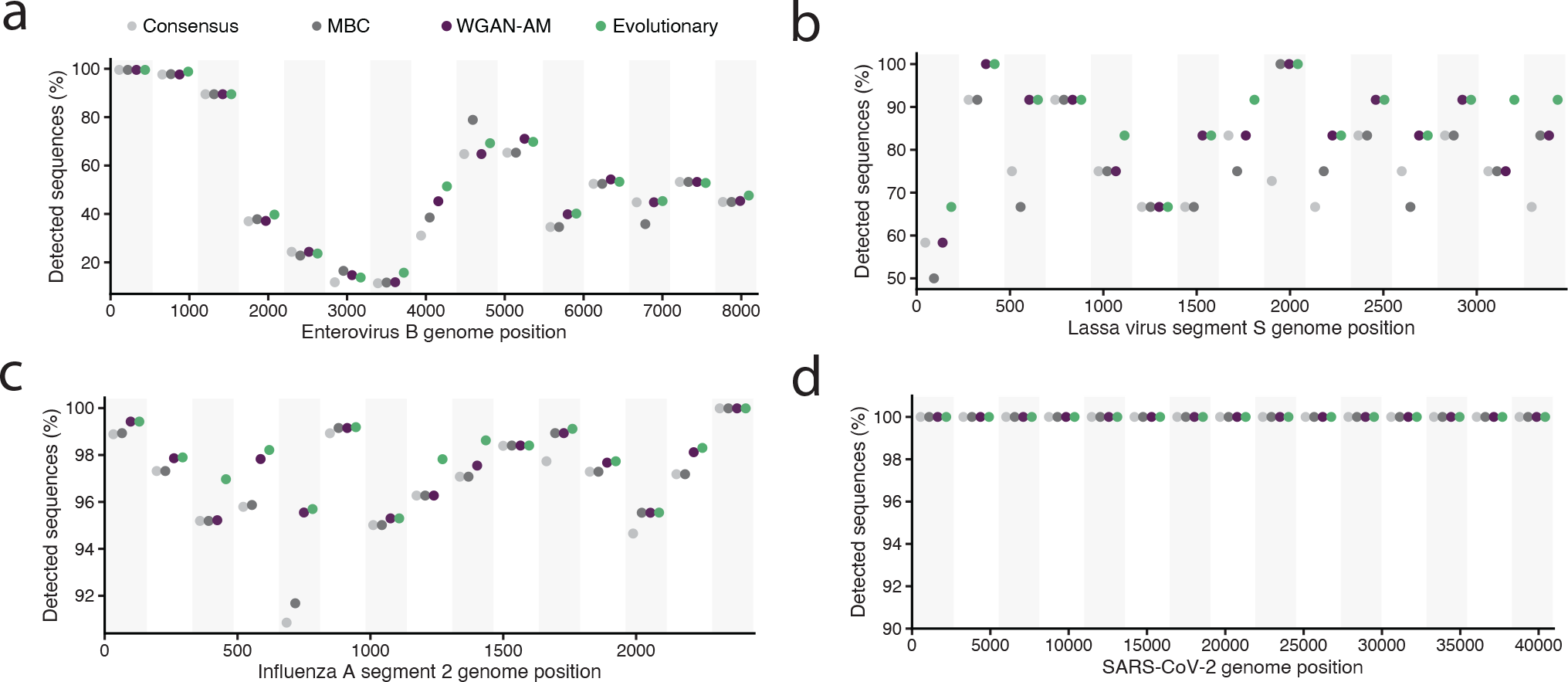
Coverage of guides designed by the MEAs and baseline methods. Proportion of genomes predicted to be detected by the guides designed with MEAs (purple and green) and baseline methods (light and dark gray) for **(a)** enterovirus B, **(b)** Lassa virus segment S, **(c)** influenza A virus segment 2, and **(d)** SARS-CoV-2. A guide is considered to detect a target if it meets the criteria described in the Methods.

**Supplementary Figure 4.**
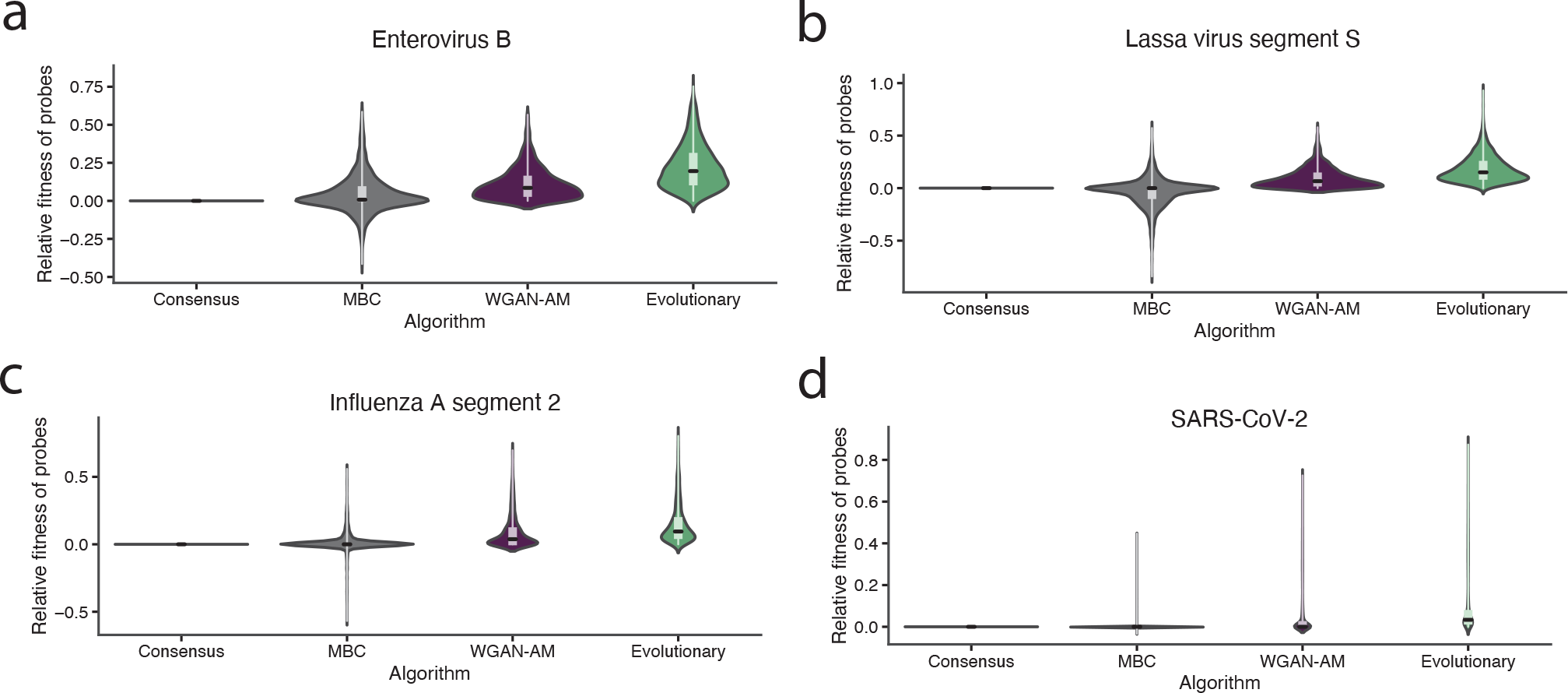
Relative fitness of guides designed by the MEAs and baseline methods. The relative fitness of the guides designed by MBC, WGAN-AM, and evolutionary algorithms at sites in the **(a)** enterovirus B, **(b)** Lassa virus segment S, **(c)** influenza A virus segment 2, and **(d)** SARS-CoV-2 genomes. The relative fitness is the difference between the fitness of the labeled algorithm’s guide and the fitness of the consensus sequence guide (by definition, consensus guides have a fitness of 0, so positive plotted values indicate fitnesses exceeding the consensus guide’s fitness). The distribution is across targetable genomic sites (see Methods).

**Supplementary Figure 5.**
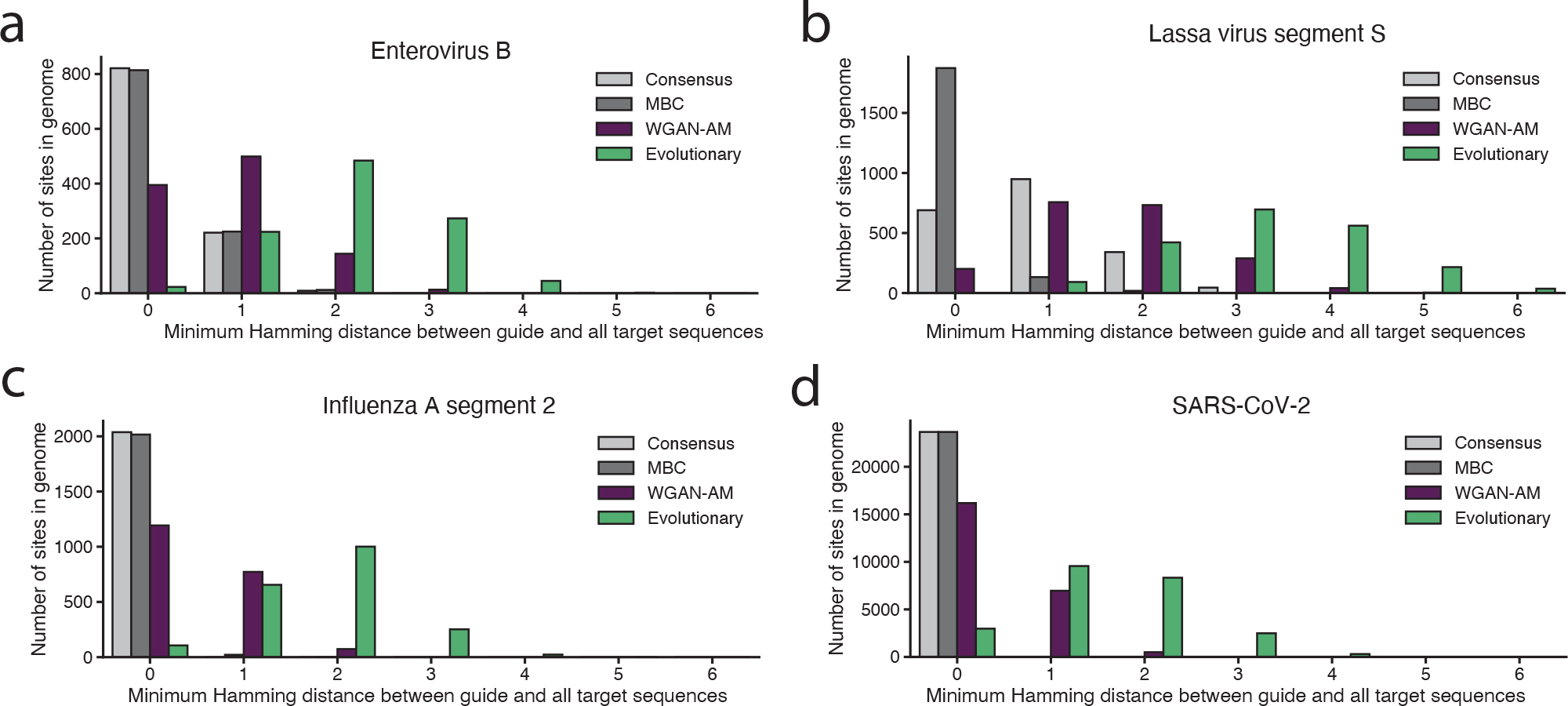
Divergence of the MEA-designed guides from observed sequences. Histograms representing the minimum Hamming distance between the guides designed by the MEA and baseline algorithms compared to all target sequences at a given genomic site—that is, the Hamming distance between a guide and the target sequence most similar to that guide. Shown for **(a)** enterovirus B, **(b)** Lassa virus segment S, **(c)** influenza A virus segment 2, and **(d)** SARS-CoV-2. The WGAN-AM (purple) and, especially, the evolutionary (green) algorithms tend to produce guides that are more dissimilar to any of the target sequences (that is, more artificial) than the baseline algorithms (light and dark gray).

**Supplementary Figure 6.**
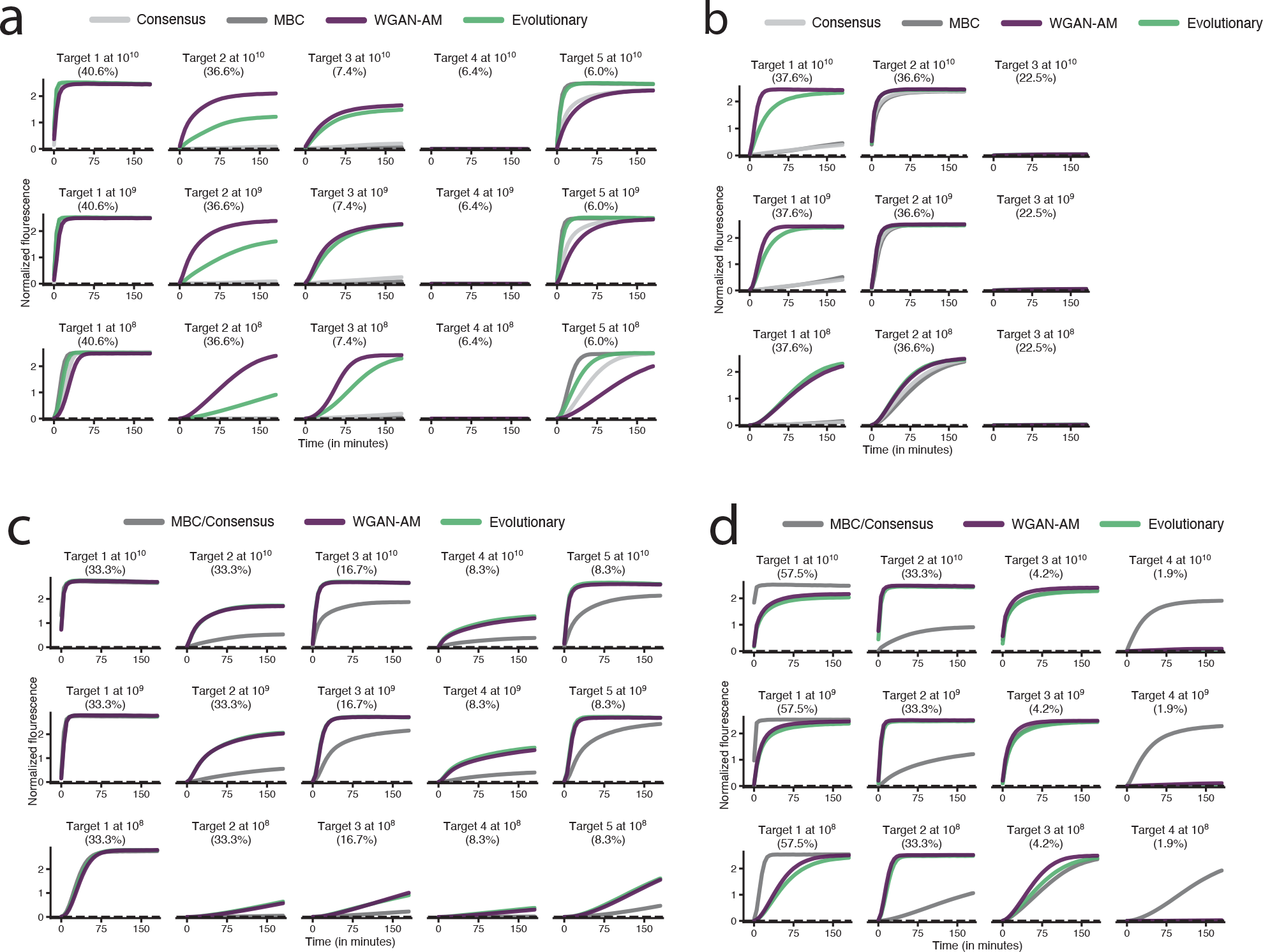
Kinetic curves for multi-target detection tasks. Normalized fluorescence over the course of the reaction for the guides targeting the genomic sites whose heatmaps are shown in Fig. 2. Parentheticals indicate the percentage of all genomes represented by the target and concentrations of targets are indicated above each plot with units of copies/μL. **(a)** Kinetic curves for a site in dengue virus (heatmap in Fig. 2e). **(b)** Kinetic curves for a site in enterovirus B (heatmap in Fig. 2f). **(c)** Kinetic curves for a site in Lassa virus segment S (heatmap in Fig. 2g). **(d)** Kinetic curves for a site in influenza A virus segment 2 (heatmap in Fig. 2d).

**Supplementary Figure 7.**
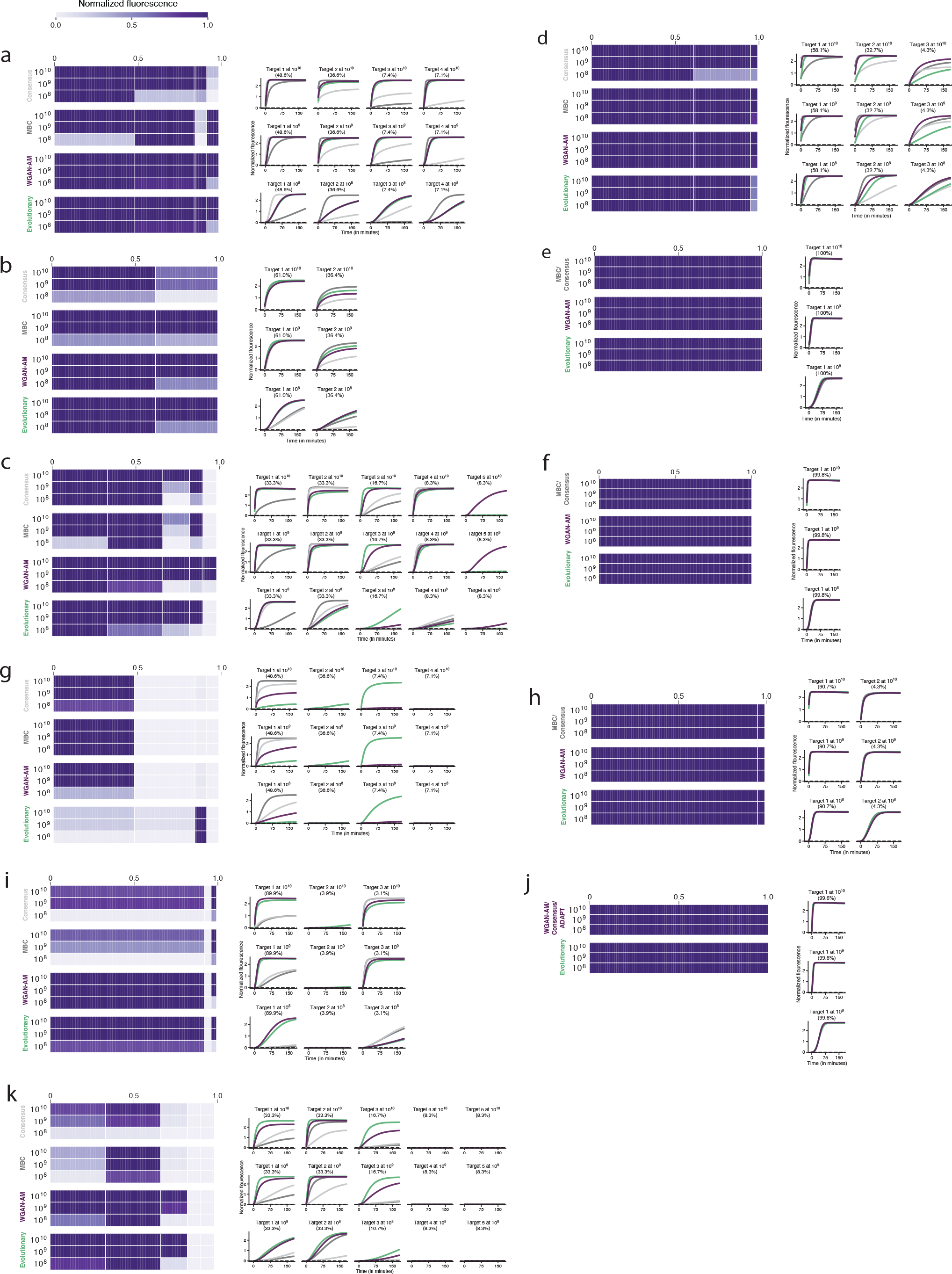
Experimental performance of multi-target detection guides. Each panel in this figure shows the experimental performance of guides designed for a specific genomic site. Panels (a–f) represent genomic sites in the top quartile of relative performance, while panels (g–k) represent genomic sites in the bottom quartile of relative performance (RP; see Methods for definition). The experimental data for the other four genomic sites in the top quartile of relative performance is shown in Fig. 2 and Supplementary Fig. 6. The left side of each panel is a heatmap representing fluorescence at 1 hour. Each column represents a target and has width proportional to the percentage of sequence diversity it represents, while each row is a concentration of the target sequence in copies/μL. The right side of each panel is a set of curves representing the normalized fluorescence of the guides against the specified targets. The color of these curves matches the color of the design method used to label each heatmap. Parentheticals indicate the percentage of all genomes represented by the target. **(a)** Site in dengue virus (top quartile of *RP*). **(b)** Site in enterovirus B (top quartile of *RP*). **(c)** Site in Lassa virus segment S (top quartile of *RP*). **(d)** Site in influenza A virus segment 2 (top quartile of *RP*). **(e)** Site in SARS-CoV-2 (top quartile of *RP*). **(f)** Site in SARS-CoV-2 (top quartile of *RP*). **(g)** Site in dengue virus (bottom quartile of *RP*). **(h)** Site in influenza A (bottom quartile of *RP*). **(i)** Site in enterovirus B (bottom quartile of *RP*). **(j)** Site in SARS-CoV-2 (bottom quartile of *RP*). **(k)** Site in Lassa virus (bottom quartile of *RP*).

**Supplementary Figure 8.**
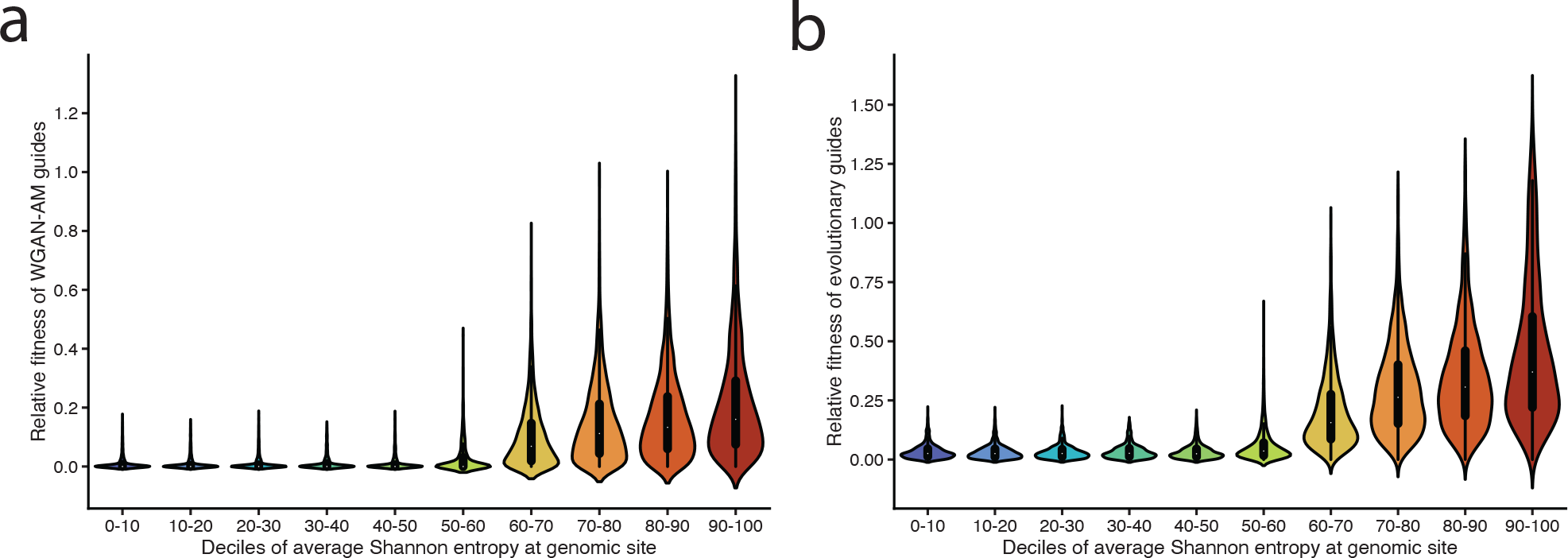
Improvement of the MEAs over baseline methods when targeting variably diverse genomic sites. Relative improvement of MEA-designed guides over the consensus guides grouped by the Shannon entropies of the genomic sites. The distribution is across all genomic sites in the 5 viral species considered for the multi-target detection objective (dengue virus, influenza A virus, enterovirus B, Lassa virus, and SARS-CoV-2). Relative fitness is defined as the fitness of the guide subtracted by the fitness of the consensus guide at that genomic site. By definition, if a guide has a relative fitness greater than zero, it is more fit than the consensus guide. The dots in the center of each violin represent the mean relative fitness for that decile. **(a)** Distribution of relative fitness for guides designed by the WGAN-AM algorithm. **(b)** Distribution of relative fitness for guides designed by the evolutionary algorithm.

**Supplementary Figure 9.**
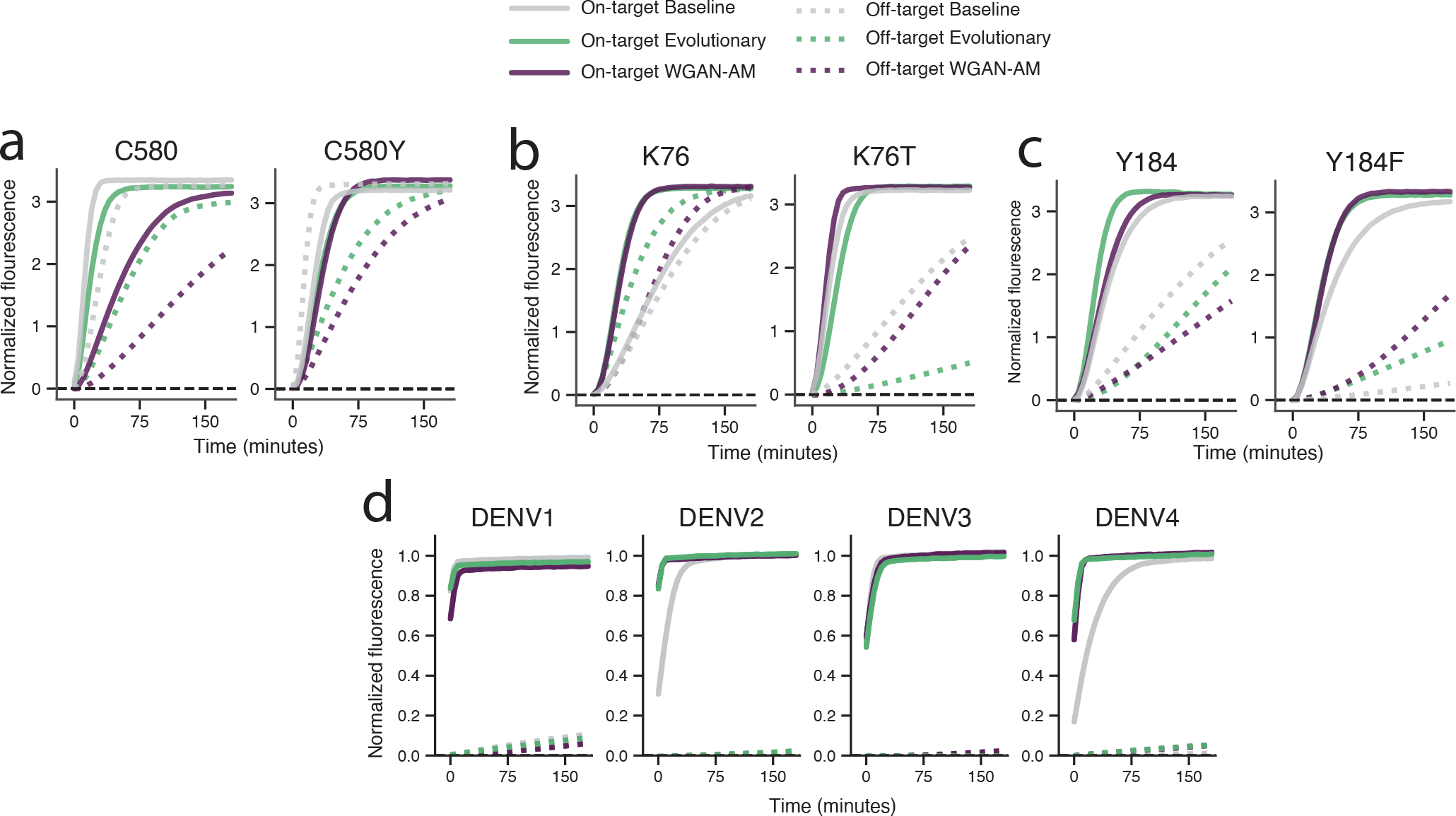
Experimental performance of guides designed for SNP and variant identification tasks. Normalized fluorescence over time is shown for **(a)** C580Y in Pfk13, **(b)** K76T in Pfcrt, **(c)** Y184F in Pfmdr1, and **(d)** dengue virus serotypes 1-4. The title of each plot indicates the target the guides were designed to detect as the on-target. On-target solid curves represent the fluorescence of the guide for its on-target and the dotted curves represent the fluorescence of the guide against its off-target. In (d), there are multiple off-targets for each discrimination task, so the dotted off-target curve shows the maximum fluorescence of the guide across the off-targets computed at each time point (e.g., the off-target curves for DENV1 represent the maximal fluorescence across the DENV2, DENV3, and DENV4 targets at each time point in the reaction). All targets were present at a concentration of 10^8^ copies/μL.

**Supplementary Figure 10.**
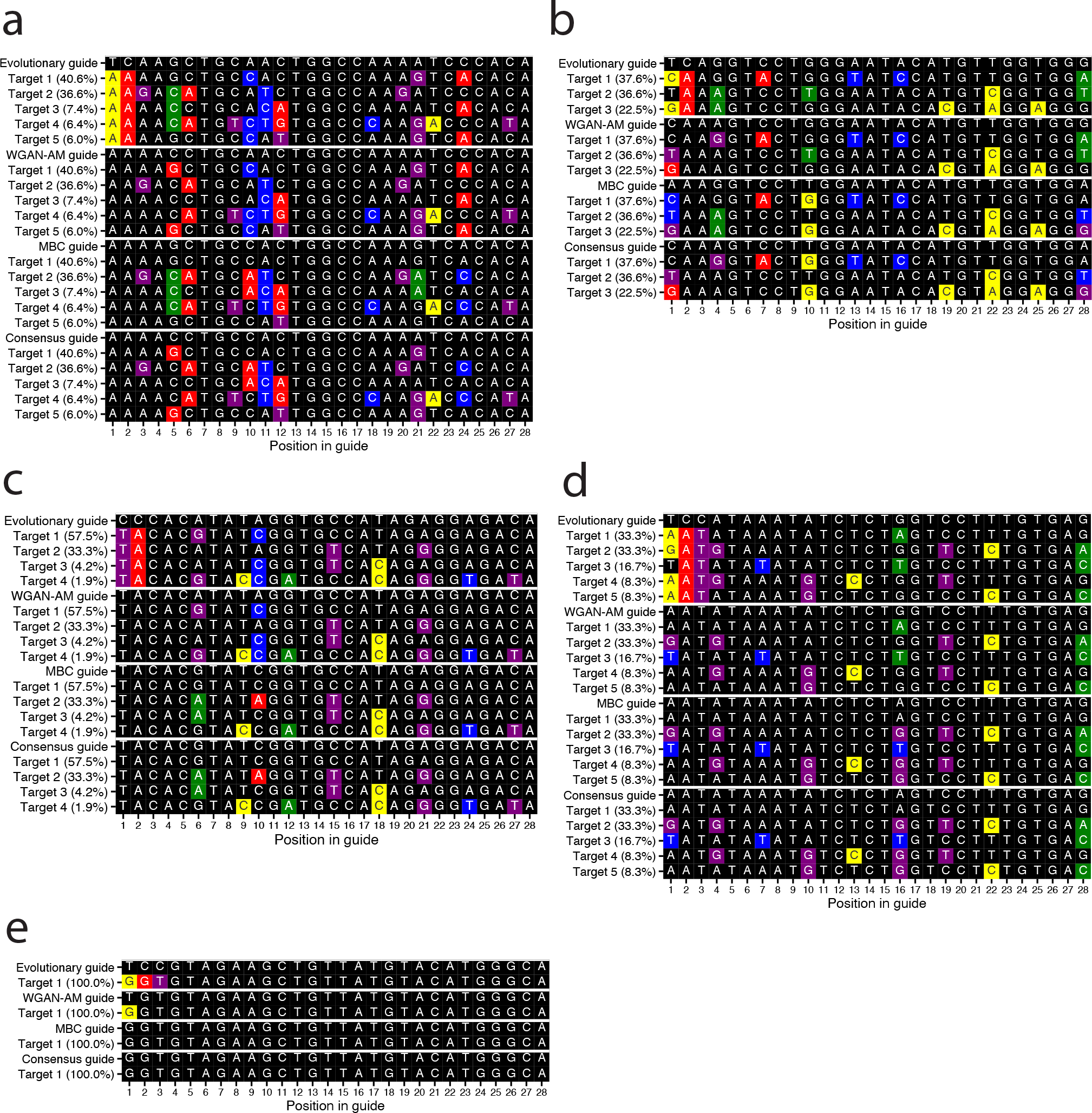
Visualization of guide-target mismatches in guides designed for the multi-target detection objective. Each panel shows guide sequences and representative target sequences for one experimentally-tested genomic site in each of the five viral pathogens considered. The guides are reverse-complemented, from their sequence in a crRNA, to be in the same frame as the target sequences, and both the guides and targets are shown as DNA sequences (with T replacing U). The black blocks indicate that there is no mismatch between the guide and target, while the blue, green, yellow, and red blocks represent A-B, G-H, T-V, and C-D mismatches respectively. The blocks in purple represent G-U wobble RNA base pairing. Parentheticals next to target names indicate the percent of sequence variation represented by that target. **(a)** A site in dengue virus. The baseline guides have mismatches with target 2 at positions 10 and 11 and also have mismatches with target 3 at positions 10, 11, and 12. The MEAs mutate the guide at position 10 to remove a mismatch at this position with targets 2 and 3, but introduce a mismatch at this position with targets 1, 4, and 5. Experimentally, the baseline guides have nearly no fluorescence on targets 2 and 3; in contrast, the MEA-designed guides achieve strong fluorescence on targets 2 and 3, while retaining similar performance on targets 1, 4, and 5 (Fig. 2e). Thus, the MEA-designed guides can detect nearly 44% more sequence diversity. (Legend continued on the subsequent page.) **(b)** A site in enterovirus B. Here, the guides designed by the baseline approaches have three mismatches in close proximity of one another (at positions 10, 13, and 16) with target 1, the target most representative of sequence variation. The MEAs mutate the nucleotide at position 10 and avoid the deleterious effect of this triple mismatch, but do introduce a mismatch with target 2 at this position. The baseline guides achieve nearly no fluorescence on target 1, while the MEA-designed guides are highly active on target 1, and still retain robust activity on target 2 (Fig. 2f). **(c)** A site in influenza A virus segment 2. The baseline guides have mismatches with target 2 at positions 6 and 10. The MEAs eliminate the position 6 mismatch against target 2 while not introducing mismatches to other targets (e.g., target 1) by employing G-U wobble base pairing between the RNAs. The MEAs also mutate the nucleotide at position 10 to remove the mismatch against target 2, but by doing this, they introduce a mismatch at position 10 against target 1. These sequence changes enable the MEA-designed guides to nearly triple the fluorescence against target 2 as the baseline guides. However, because they do introduce a mismatch to target 1, they have slightly lower fluorescence at the initial timepoints against this target but still achieve saturating fluorescence by the later timepoints of the reaction (Fig. 2d). **(d)** A site in Lassa virus segment S. At this site, MEA-designed guides achieve substantially greater fluorescence on both targets 2 and 4 (Fig. 2g). From the pattern of guide-target mismatches, it is not clear why the MEA-designed guides perform better. As one hypothesis, the close proximity of the instances of G-U wobble base pairing at positions 16 and 19 of the baseline guides might be deleterious to activity; here, the MEAs might be exploiting features of Cas13a targeting that are not currently well-characterized. **(e)** A site in SARS-CoV-2. Previous work (Fig. 2e in ref. 22) has shown that a T-V guide-target mismatch at position 1 is strongly associated with increased guide activity. Both the WGAN-AM and evolutionary algorithms introduce this mismatch. Experimentally, the evolutionary and WGAN-AM guides both achieve slightly greater fluorescence than the baseline (MBC and consensus) guides, likely due to their position 1 mismatch (Supplementary Fig. 7e).

**Supplementary Figure 11.**
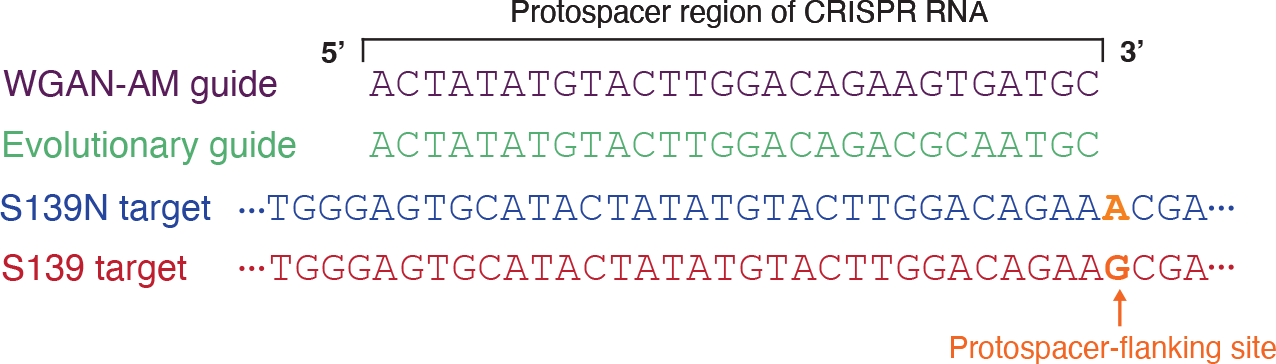
MEAs exploit Cas13a’s protospacer context preferences to design optimal variant identification guides. When targeting a SNP from a non-G nucleotide a G nucleotide, the MEAs often place the G nucleotide in the off-target at the PFS for optimal SNP discrimination. In this schematized example, the guides were designed to achieve high activity on the S139N target and low activity on the S139 target. Both the WGAN-AM and evolutionary algorithm positioned the guides such that the G nucleotide in the S139 off-target sequence is located at the PFS, thus lowering off-target activity, as is experimentally observed (Fig. 3e).

**Supplementary Figure 12.**
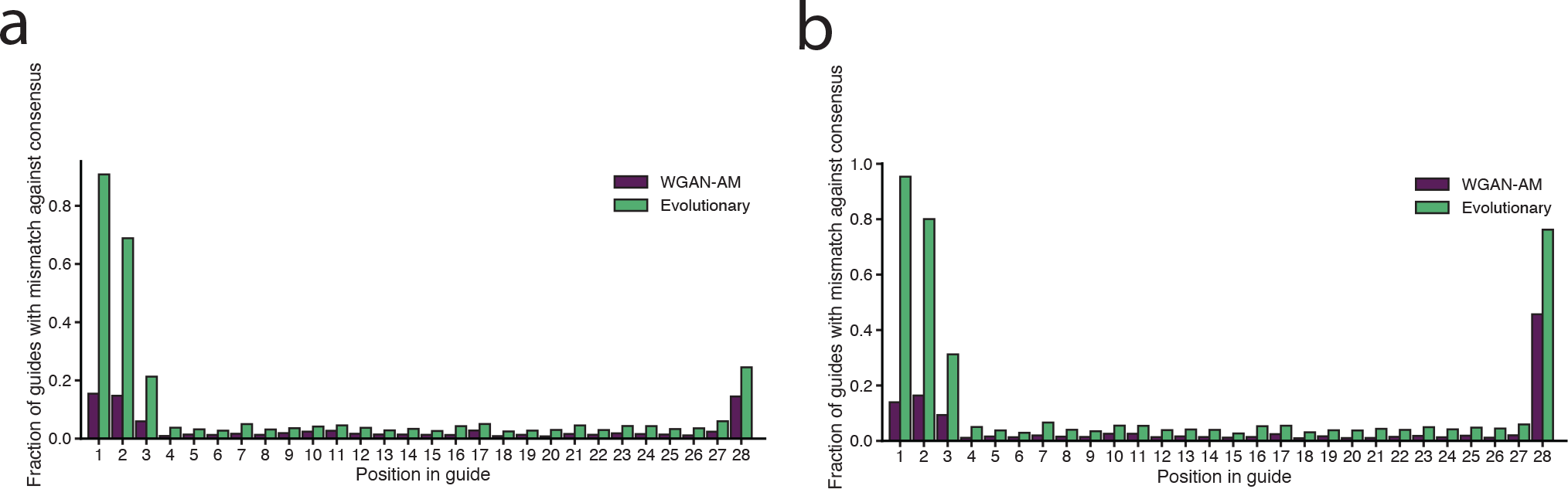
Divergence of MEA-designed guides from the consensus sequence. Fraction of MEA-designed guides for the multi-target detection objective that have a mismatch against the consensus genome sequence, at each position in the guide-target pairing. **(a)** Fractions taken across all genomic sites from all of the five viral pathogens considered (dengue virus, influenza A virus, enterovirus B, Lassa virus, and SARS-CoV-2). **(b)** Fractions taken across only the genomic sites with a G nucleotide at the PFS, again across all of the five viral pathogens considered. On this subset of sites, the MEA-designed guides are relatively likely to have a terminal mismatch in the protospacer, suggesting the benefit of such a mismatch when there is a G nucleotide at the PFS.

**Supplementary Figure 13.**
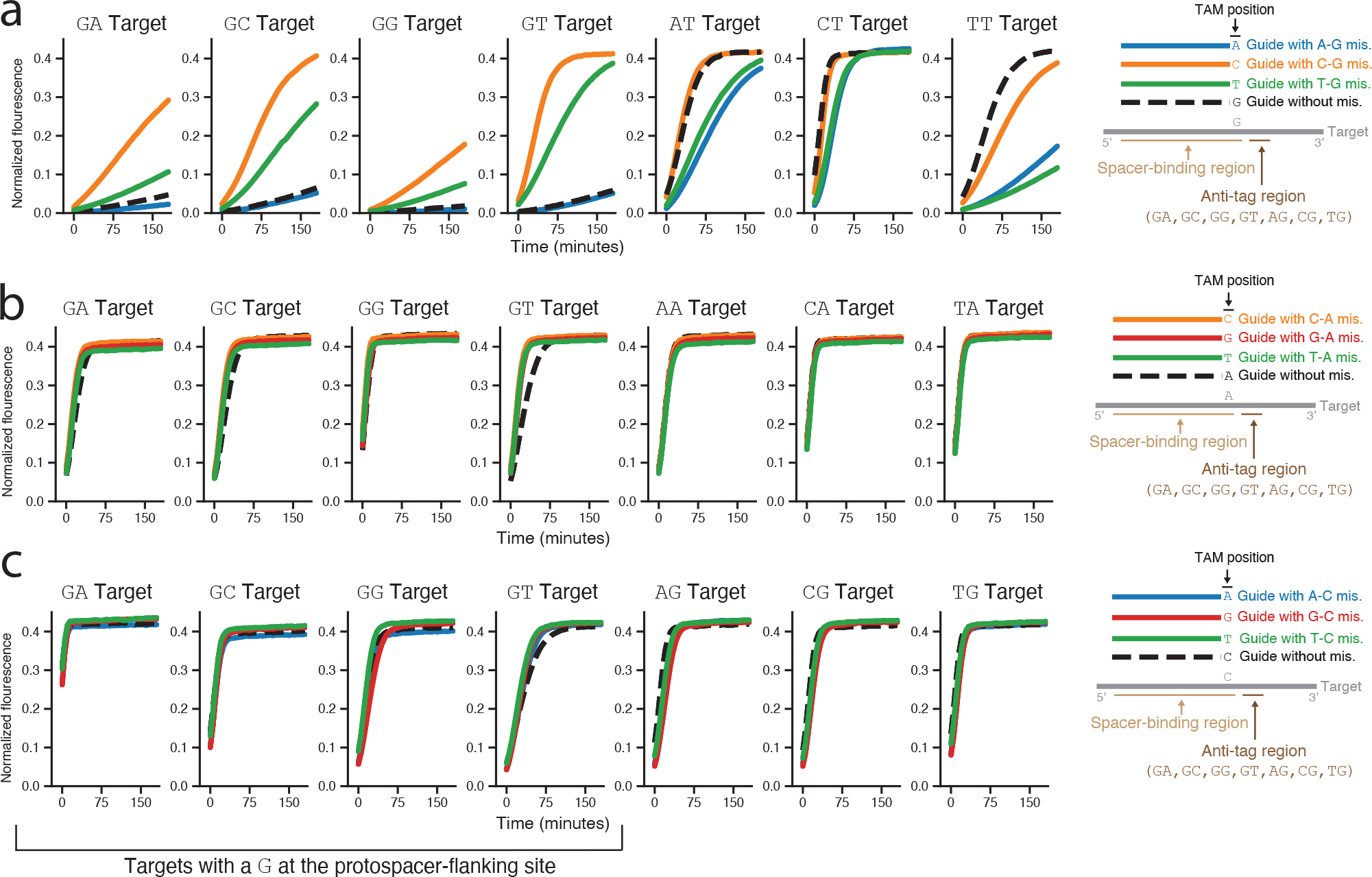
Evaluation of the ability of the tag-adjacent mismatch to rescue guide activity for different target sets. Normalized fluorescence curves are shown for the target sets where the tag-adjacent position (TAM position) in the target is a **(a)** G nucleotide, **(b)** A nucleotide, or **(c)** C nucleotide. All guides are identical to the targets at positions 1-27 in the protospacer. The colored lines represent guides with a tag-adjacent mismatch (at position 28), while the black dashed lines represent guides without a mismatch to the target. The dinucleotide above each plot indicates the first two nucleotides of that target’s anti-tag region, and the guide sequences represented in the schematic are reverse-complemented compared to the spacer sequence in the CRISPR RNA. The same curves for the target set with a T nucleotide at the tag-adjacent position are shown in Fig. 3i.

**Supplementary Figure 14.**
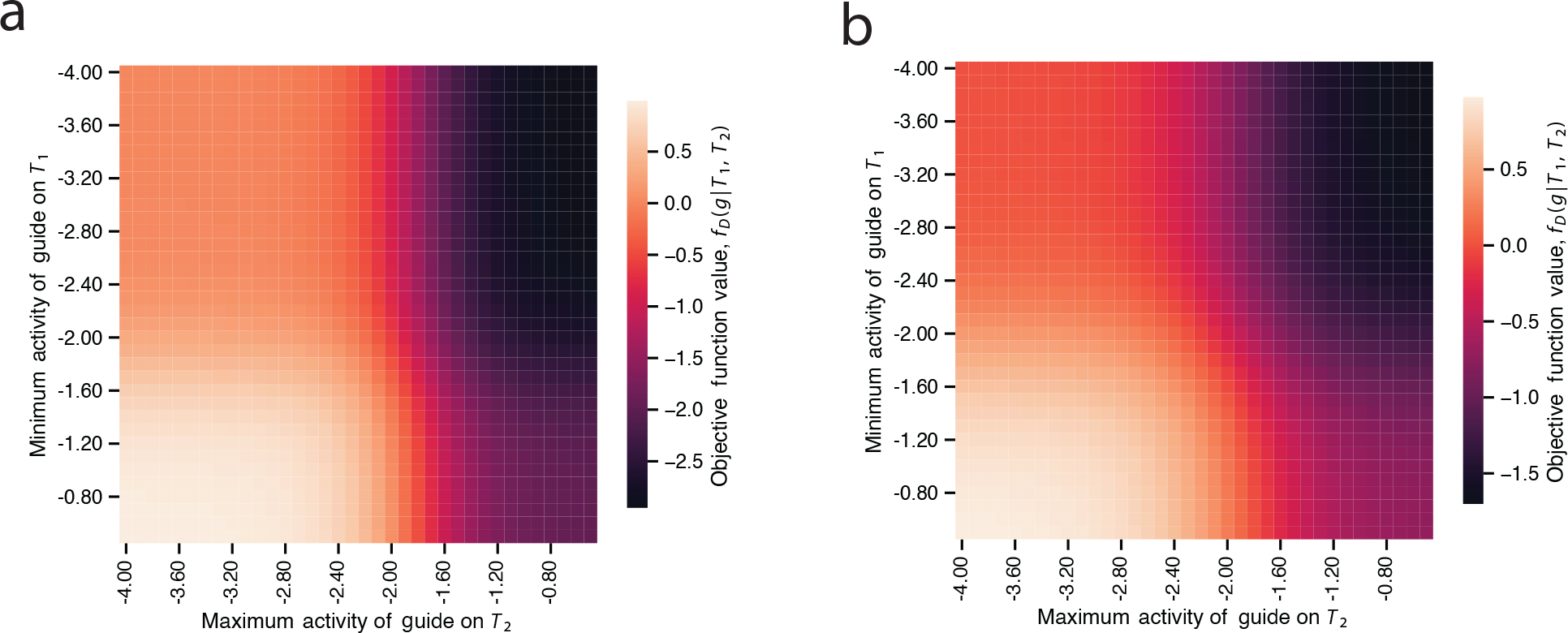
Visualization of the fitness function for the variant identification objective. Colors represent the value of the variant identification fitness function according to on-target (*T*_1_) and off-target (*T*_2_) activities (see Methods for the function). A maximally-fit guide *g* has a low maximum activity across the off-target set *T*_2_, 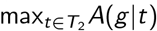, and a high minimum activity across the on-target set *T*_1_, 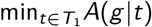. **(a)** Using hyperparameters determined by the WGAN-AM random search. **(b)** Using hyperparameters determined by the evolutionary algorithm random search.

**Supplementary Figure 15.**
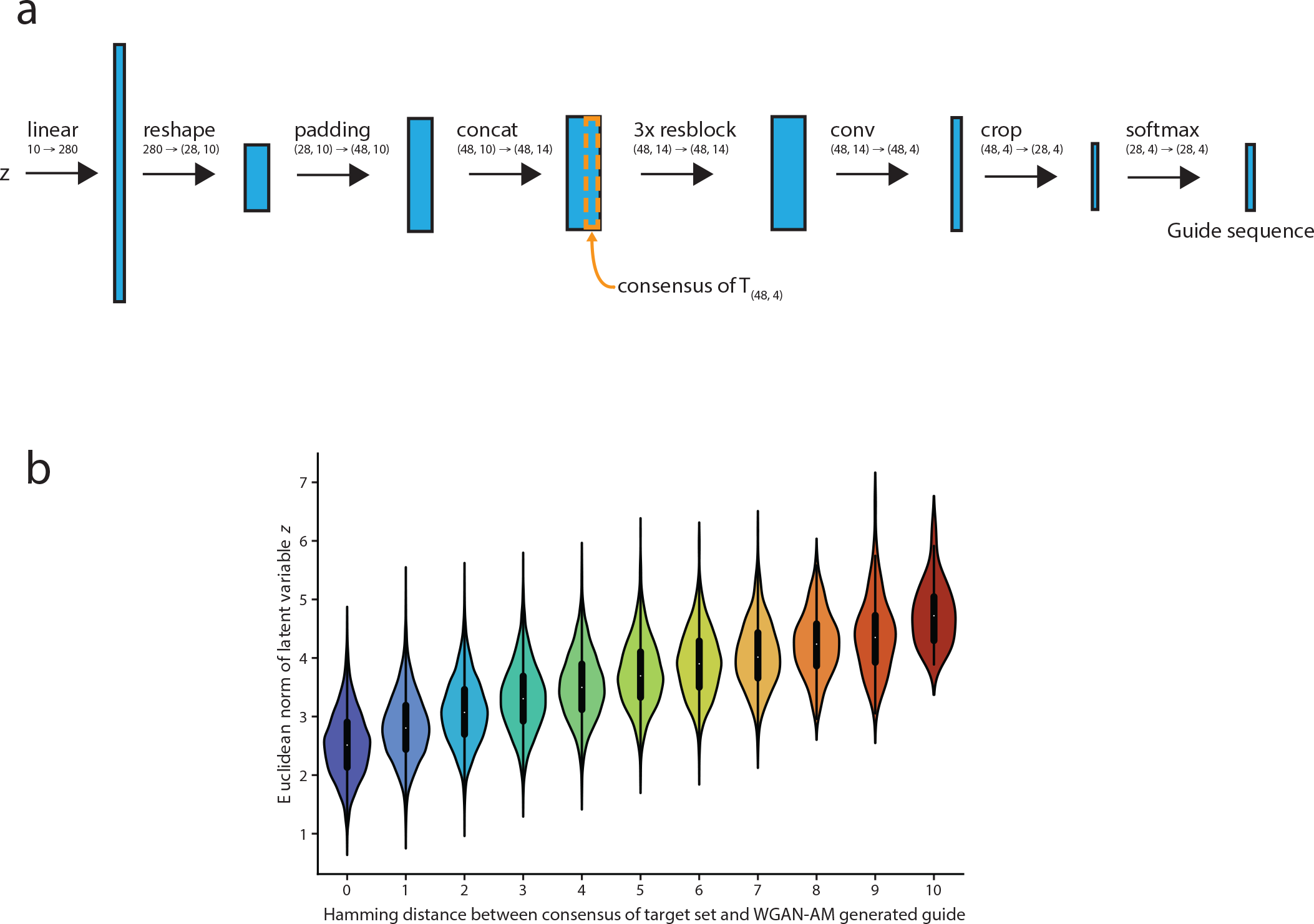
Properties of WGAN-AM algorithm. **(a)** Architecture of generative model of Wasserstein generative adversarial network (WGAN) for generating guides, inspired by that of ref. 25. The input into the generative model, the latent vector *z*, is a 10-dimensional vector sampled from the standard normal distribution. *z* is upsampled through a linear layer and reshaped and padded to create a 48 by 10 matrix. In the concat operation, the one-hot-encoded consensus sequence of the target set *T* (with dimensions 48 by 4) is concatenated with the output of the previous layer to create a 48 by 14 matrix. This matrix is passed through three residual blocks. Each residual block (resblock) consists of two layers, with each layer containing a 1D convolutional layer with 14 filters of stride 1 and width 3. Next, in the conv operation, a convolutional layer with 4 filters of width 1 and stride 1 is applied to the output of the resblock to create 4-channel encoded guide sequence. Finally, the matrix is cropped to remove the 10 by 4 context on each side of the guide protospacer sequence, and the resulting 28 by 4 matrix is passed through a softmax layer so that each element in the matrix represents the probability of having a certain base at that position. The guide sequence contains the base at each position with the greatest probability. **(b)** Relationship between the Euclidean norm of the latent variable input to the WGAN’s generator network and the number of mutations introduced into the guide relative to the consensus sequence. All guides with a Hamming distance ≤ 10 are shown, and the computational experiment was performed as described in the Methods.

## References

[1] Sinai, S. & Kelsic, E. D. A primer on model-guided exploration of fitness landscapes for biological sequence design. arXiv (2020). 2010.10614.

[2] Sinai, S. et al. Adalead: A simple and robust adaptive greedy search algorithm for sequence design. arXiv preprint (2020).

[3] Brookes, D., Park, H. & Listgarten, J. Conditioning by adaptive sampling for robust design. In Chaudhuri, K. & Salakhutdinov, R. (eds.) Proceedings of the 36th International Conference on Machine Learning, vol. 97 of Proceedings of Machine Learning Research, 773–782 (PMLR, 2019).

[4] Linder, J., Bogard, N., Rosenberg, A. B. & Seelig, G. A generative neural network for maximizing fitness and diversity of synthetic DNA and protein sequences. Cell Syst 11, 49–62.e16 (2020).

[5] Valeri, J. A. et al. Sequence-to-function deep learning frameworks for engineered riboregulators. Nat. Commun. 11, 5058 (2020).

[6] Gupta, A. & Zou, J. Feedback GAN for DNA optimizes protein functions. Nature Machine Intelligence 1, 105–111 (2019).

[7] Repecka, D. et al. Expanding functional protein sequence spaces using generative adversarial networks. Nature Machine Intelligence 3, 324–333 (2021).

[8] Shin, J.-E. et al. Protein design and variant prediction using autoregressive generative models. Nat. Commun. 12, 2403 (2021).

[9] Watson, J. L. et al. Broadly applicable and accurate protein design by integrating structure prediction networks and diffusion generative models. bioRxiv 2022.12.09.519842 (2022).

[10] Strokach, A. & Kim, P. M. Deep generative modeling for protein design. Curr. Opin. Struct. Biol. 72, 226–236 (2022).

[11] Madani, A. et al. Large language models generate functional protein sequences across diverse families. Nature Biotechnology (2023).

[12] Shanehsazzadeh, A. et al. Unlocking de novo antibody design with generative artificial intelligence. bioRxiv (2023). URL https://www.biorxiv.org/content/early/2023/01/09/2023.01.08.523187. https://www.biorxiv.org/content/early/2023/01/09/2023.01.08.523187.full.pdf.

[13] Doench, J. G. et al. Optimized sgRNA design to maximize activity and minimize off-target effects of CRISPR-Cas9. Nat. Biotechnol. 34, 184–191 (2016).

[14] Listgarten, J. et al. Prediction of off-target activities for the end-to-end design of CRISPR guide RNAs. Nat Biomed Eng 2, 38–47 (2018).

[15] Chuai, G. et al. DeepCRISPR: optimized CRISPR guide RNA design by deep learning. Genome Biol. 19, 80 (2018).

[16] Moreno-Mateos, M. A. et al. CRISPRscan: designing highly efficient sgRNAs for CRISPR-Cas9 targeting in vivo. Nat. Methods 12, 982–988 (2015).

[17] Kim, H. K. et al. Deep learning improves prediction of CRISPR-Cpf1 guide RNA activity. Nat. Biotechnol. 36, 239–241 (2018).

[18] Chari, R., Yeo, N. C., Chavez, A. & Church, G. M. sgRNA scorer 2.0: A Species-Independent model to predict CRISPR/Cas9 activity. ACS Synth. Biol. 6, 902–904 (2017).

[19] Montague, T. G., Cruz, J. M., Gagnon, J. A., Church, G. M. & Valen, E. CHOPCHOP: a CRISPR/Cas9 and TALEN web tool for genome editing. Nucleic Acids Res. 42, W401–7 (2014).

[20] Heigwer, F., Kerr, G. & Boutros, M. E-CRISP: fast CRISPR target site identification. Nat. Methods 11, 122–123 (2014).

[21] Liu, G., Zhang, Y. & Zhang, T. Computational approaches for effective CRISPR guide RNA design and evaluation. Computational and Structural Biotechnology Journal 18, 35–44 (2020).

[22] Metsky, H. C. et al. Designing sensitive viral diagnostics with machine learning. Nat Biotechnology 40, 1123–1131 (2022).

[23] Zhao, D. et al. Imperfect guide-RNA (igRNA) enables CRISPR single-base editing with ABE and CBE. Nucleic Acids Res 50, 4161–4170 (2022).

[24] Gootenberg, J. S. et al. Nucleic acid detection with CRISPR-Cas13a/C2c2. Science 356, 438–442 (2017).

[25] Killoran, N., Lee, L. J., Delong, A., Duvenaud, D. & Frey, B. J. Generating and designing dna with deep generative models. arXiv (2017). 1712.06148.

[26] Yuan, L. et al. A single mutation in the prm protein of zika virus contributes to fetal microcephaly. Science 358, 933–936 (2017).

[27] Apinjoh, T. O., Ouattara, A., Titanji, V. P. K., Djimde, A. & Amambua-Ngwa, A. Genetic diversity and drug resistance surveillance of plasmodium falciparum for malaria elimination: is there an ideal tool for resource-limited sub-saharan africa? Malar. J. 18, 217 (2019).

[28] Carter, T. E. et al. Evaluation of dihydrofolate reductase and dihydropteroate synthetase genotypes that confer resistance to sulphadoxine-pyrimethamine in plasmodium falciparum in haiti (2012).

[29] Quan, H. et al. High multiple mutations of plasmodium falciparum-resistant genotypes to sulphadoxine-pyrimethamine in lagos, nigeria. Infect Dis Poverty 9, 91 (2020).

[30] Yoshida, N., Yamauchi, M., Morikawa, R., Hombhanje, F. & Mita, T. Increase in the proportion of plasmodium falciparum with kelch13 C580Y mutation and decline in pfcrt and pfmdr1 mutant alleles in papua new guinea. Malar. J. 20, 410 (2021).

[31] Hyde, J. E. Drug-resistant malaria. Trends Parasitol. 21, 494–498 (2005).

[32] Harvey, W. T. et al. SARS-CoV-2 variants, spike mutations and immune escape. Nat Rev Microbiol 19, 409–424 (2021).

[33] Tambe, A., East-Seletsky, A., Knott, G. J., Doudna, J. A. & O’Connell, M. R. RNA binding and HEPN-Nuclease activation are decoupled in CRISPR-Cas13a. Cell Rep. 24, 1025–1036 (2018).

[34] Abudayyeh, O. O. et al. RNA targeting with CRISPR-Cas13. Nature 550, 280–284 (2017).

[35] Abudayyeh, O. O. et al. C2c2 is a single-component programmable RNA-guided RNA-targeting CRISPR effector. Science 353, aaf5573 (2016).

[36] Meeske, A. J. & Marraffini, L. A. RNA guide complementarity prevents Self-Targeting in type VI CRISPR systems. Mol. Cell 71, 791–801.e3 (2018).

[37] Welch, N. L. et al. Multiplexed CRISPR-based microfluidic platform for clinical testing of respiratory viruses and identification of SARS-CoV-2 variants. Nature Medicine (2022).

[38] Jabado, O. J. et al. Greene SCPrimer: a rapid comprehensive tool for designing degenerate primers from multiple sequence alignments. Nucleic Acids Research 34, 6605–6611 (2006).

[39] Kreer, C. et al. openPrimeR for multiplex amplification of highly diverse templates. J. Immunol. Methods 480, 112752 (2020).

[40] Duitama, J. et al. PrimerHunter: a primer design tool for PCR-based virus subtype identification. Nucleic Acids Res. 37, 2483–2492 (2009).

[41] Brodin, J. et al. A multiple-alignment based primer design algorithm for genetically highly variable DNA targets. BMC Bioinformatics 14, 255 (2013).

[42] Bock, C. et al. High-content CRISPR screening. Nat Rev Methods Primers 2 (2022).

[43] Huang, X., Yang, D., Zhang, J., Xu, J. & Chen, Y. E. Recent Advances in Improving Gene-Editing Specificity through CRISPR-Cas9 Nuclease Engineering. Cells 11 (2022).

[44] Saha, K. Accounting for diversity in the design of CRISPR-based therapeutic genome editing. Nat Genet 55, 6–7 (2023).

[45] Goodfellow, I. J. et al. Generative adversarial networks. arXiv (2014). 1406.2661.

[46] Mirza, M. & Osindero, S. Conditional generative adversarial nets. arXiv (2014). URL https://arxiv.org/abs/1411.1784.

[47] Gulrajani, I., Ahmed, F., Arjovsky, M., Dumoulin, V. & Courville, A. Improved training of wasserstein gans. arXiv preprint arXiv:1704.00028 (2017).

[48] Kingma, D. P. & Ba, J. Adam: A method for stochastic optimization. arXiv (2014). 1412.6980.

[49] Porto, W. F. et al. In silico optimization of a guava antimicrobial peptide enables combinatorial exploration for peptide design. Nature Communications 9 (2018).

[50] Yoshida, M. et al. Using evolutionary algorithms and machine learning to explore sequence space for the discovery of antimicrobial peptides. Chem 4, 533–543 (2018).

[51] Sinai, S. & Kelsic, E. D. A primer on model-guided exploration of fitness landscapes for biological sequence design. arXiv (2020). 2010.10614.

[52] Martín Abadi et al. TensorFlow: Large-Scale machine learning on heterogeneous systems (2015).

[53] Federhen, S. The NCBI taxonomy database. Nucleic Acids Research 40, D136–43 (2012).

[54] Katoh, K. & Standley, D. M. MAFFT multiple sequence alignment software version 7: improvements in performance and usability. Molecular Biology and Evolution 30, 772–780 (2013).

[55] Shu, Y. & McCauley, J. GISAID: Global initiative on sharing all influenza data - from vision to reality. Euro surveillance: bulletin Europeen sur les maladies transmissibles = European communicable disease bulletin 22 (2017).

[56] Pickett, B. E. et al. ViPR: An open bioinformatics database and analysis resource for virology research. Nucleic Acids Research 40 (2011).

[57] Hodcroft, E. B. CoVariants: SARS-CoV-2 mutations and variants of interest. https://covariants.org/.

[58] Mullen, J. L. et al. outbreak.info. https://outbreak.info/.

[59] Kellner, M. J., Koob, J. G., Gootenberg, J. S., Abudayyeh, O. O. & Zhang, F. SHERLOCK: nucleic acid detection with CRISPR nucleases. Nat. Protoc. 14, 2986–3012 (2019).

